# *In-situ* glial cell-surface proteomics identifies pro-longevity factors in *Drosophila*

**DOI:** 10.1101/2025.09.26.678810

**Authors:** Madeline P. Marques, Bo Sun, Ye-Jin Park, Tyler Jackson, Tzu-Chiao Lu, Yanyan Qi, Erin Harrison, Miranda C. Wang, Omar Moussa Pasha, Amogh Varanasi, Dominique Kiki Carey, D.R. Mani, Jonathan Zirin, Mujeeb Qadiri, Yanhui Hu, Kartik Venkatachalam, Norbert Perrimon, Steven A. Carr, Namrata D. Udeshi, Liqun Luo, Jiefu Li, Hongjie Li

**Author notes:** Corresponding author (H.L.).

## Abstract

Much focus has shifted towards understanding how glial dysfunction contributes to age-related neurodegeneration due to the critical roles glial cells play in maintaining healthy brain function. Cell-cell interactions, which are largely mediated by cell-surface proteins, control many critical aspects of development and physiology; as such, dysregulation of glial cell-surface proteins in particular is hypothesized to play an important role in age-related neurodegeneration. However, it remains technically difficult to profile glial cell-surface proteins in intact brains. Here, we applied a cell-surface proteomic profiling method to glial cells from intact brains in *Drosophila,* which enabled us to fully profile cell-surface proteomes *in-situ*, preserving native cell-cell interactions that would otherwise be omitted using traditional proteomics methods. Applying this platform to young and old flies, we investigated how glial cell-surface proteomes change during aging. We identified candidate genes predicted to be involved in brain aging, including several associated with neural development and synapse wiring molecules not previously thought to be particularly active in glia. Through a functional genetic screen, we identified one surface protein, DIP-β, which is down-regulated in old flies and can increase fly lifespan when overexpressed in adult glial cells. We further performed whole-head single-nucleus RNA-seq and revealed that DIP-β overexpression mainly impacts glial and fat cells. We also found that glial DIP-β overexpression was associated with improved cell-cell communication, which may contribute to the observed lifespan extension. Our study is the first to apply *in-situ* cell-surface proteomics to glial cells in *Drosophila*, and to identify DIP-β as a potential glial regulator of brain aging.

## Introduction

During aging, several changes occur in the brain that render it more vulnerable to a variety of age-related neurodegenerative diseases, including Alzheimer’s Disease and Parkinson’s Disease (Mattson and Arumugam, 2018). These changes include impaired neuroplasticity and resilience, aberrant neuronal network activity, disruption of homeostasis, inflammation, etc. Aging, defined as a time-related decline in physiological functions (Ebeling et al., 2021; Dorsey and Bloem, 2024), is considered the strongest risk factor for these conditions. While these neurodegenerative diseases share many phenotypic similarities with normal brain aging, we still do not fully understand the mechanisms underlying brain aging, nor how these mechanisms deviate in each of these unique pathologies (Mattson and Arumugam, 2018; Iijima et al., 2004; Drachman, 2006; Moqri et al., 2023). By characterizing the functions and mechanisms responsible for brain aging and establishing a foundational understanding of how “normal” age-related neurodegeneration occurs, we will be far better equipped to develop more targeted interventions moving forward.

Recently, much focus has shifted towards understanding how glial dysfunction may contribute to age-related neurodegeneration. This is largely due to the many crucial roles glial cells play in maintaining healthy brain function, including moderating cell-cell interactions, maintaining homeostasis, facilitating immune responses, etc. (Barres, 2008; Freeman, 2015; Kremer et al., 2017). Cell-cell interactions, which are largely mediated by cell-surface proteins, control many critical aspects of development and physiology (Sperry, 1963; Malenka and Bear, 2004; Jan and Jan, 2010; Zipursky and Sanes, 2010; Kolodkin and Tessier-Lavigne, 2011; Yeh et al., 2017; Li et al., 2020; McAlpine et al., 2021). As dysregulation of glia-neuron interactions is considered a hallmark of normal brain aging (Barres, 2008), glial cell-surface proteins are hypothesized to play important roles in maintaining healthy brain function during aging (Kremer et al., 2017). However, fully characterizing the cell-surface proteomes of glial cells has proven challenging using traditional proteomics methods, such as biochemical fractionation (Cordwell and Thingholm, 2010; Li et al., 2020). This is because biochemical fractionation does not allow for cell-type specificity, includes various contaminants (mitochondrial, ER, and Golgi), and, importantly, omits important secreted and extracellular matrix proteins that form an integral part of the cell-surface proteome (Li et al., 2020).

Recently, *in-situ* cell-surface proteomics methods were developed to profile spatiotemporally resolved neuronal cell-surface proteomes from intact brains in *Drosophila* (Li et al., 2020; Xie et al., 2022), as well as in certain mouse brain regions (Shuster et al., 2022). This technique enables complete profiling of cell-surface proteomes *in-situ*, preserving native cell-cell interactions. Ultimately, this allows investigators to form a more comprehensive understanding of how glia-neuron interactions evolve during aging (as this method effectively captures both the glial cell-surface proteins themselves, as well as any other endogenous proteins that are in very close proximity). Applying this platform to the glial cells of young and old flies, we investigated how glial cell-surface proteomes change as flies age. We identified several candidate proteins predicted to be involved in normal brain aging, including several neural development and synapse wiring molecules not previously thought to be particularly active in glia or in aging (Szklarczyk et al., 2023). We found that one synapse wiring molecule in particular, DIP-β, was associated with significant lifespan increases when overexpressed in adult glial cells (using the *glial-GeneSwitch* conditional driver). Our single-nucleus RNA-seq data suggest that glial DIP-β overexpression improves cell-cell communication, including glia-neuron and glia-fat cell interactions, contributing to lifespan extension.

Our study is the first to apply *in-situ* cell-surface proteomics to glial cells in *Drosophila*, and to fully profile and characterize the glial cell-surface proteomes of aging-model (i.e., young and old) flies. Additionally, while DIP-β’s role in precise neural circuit assembly during development has been studied extensively (Carrillo et al., 2015; Cosmanescu et al., 2018; Sanes and Zipursky, 2020; Sergeeva et al., 2020; Wang et al., 2022; Ma et al., 2023), ours is the first to identify DIP-β as a potential regulator of healthy brain function and synapse maintenance during normal brain aging. Combined, these results further demonstrate the power of *in-situ* cell-surface proteomics to identify new molecules involved in the maintenance of healthy brain function during aging.

## Results

### Glial cell-surface proteomes from young and old fly brains

To fully profile the cell-surface proteomes of glial cells in intact fly brains, we used a modified peroxidase-mediated proximity-based biotinylation procedure (Li et al., 2020; Loh et al., 2016; Xie at al., 2022). To accomplish cell-type and cell-surface specificity, we employed the glia-specific driver *Repo-GAL4* to drive transgenic expression of horseradish peroxidase (HRP) fused to the N-terminus of the transmembrane protein rat CD2 (*UAS-HRP-CD2*). This simultaneously targets HRP-CD2 to the extracellular side of the plasma membrane, while restricting HRP-CD2 expression to glial cells (Larsen et al., 2003; Watts et al., 2004; Li et al., 2020). Next, we used the biotin-xx-phenol (BxxP) substrate, which is unable to cross the plasma membrane (Loh et al., 2016; Li et al., 2020; Durojaye, 2021; Renuse et al., 2020). Combined with the glial cell-surface-targeting of HRP-CD2, this dual-gate approach is designed to enhance the specificity toward glial cell-surface proteins. In the presence of hydrogen peroxide (H2O2), HRP forms a reactive complex (HRP-H2O2), which converts BxxP into short-lived phenoxyl radicals that rapidly biotinylate endogenous proteins that are in very close proximity (**Fig. 1A**). Because the half-life of these phenoxyl radicals is extremely short (less than 1 millisecond) and the reaction requires only a few minutes to complete, phenoxyl radicals can only label proteins within a very small radius from the peroxidase enzyme (Durojaye, 2021). Thus, this provides a snapshot of the cell-surface proteome with high temporal precision, while minimizing the potentially toxic effects of hydrogen peroxide (Li et al., 2021).

**Figure 1:**
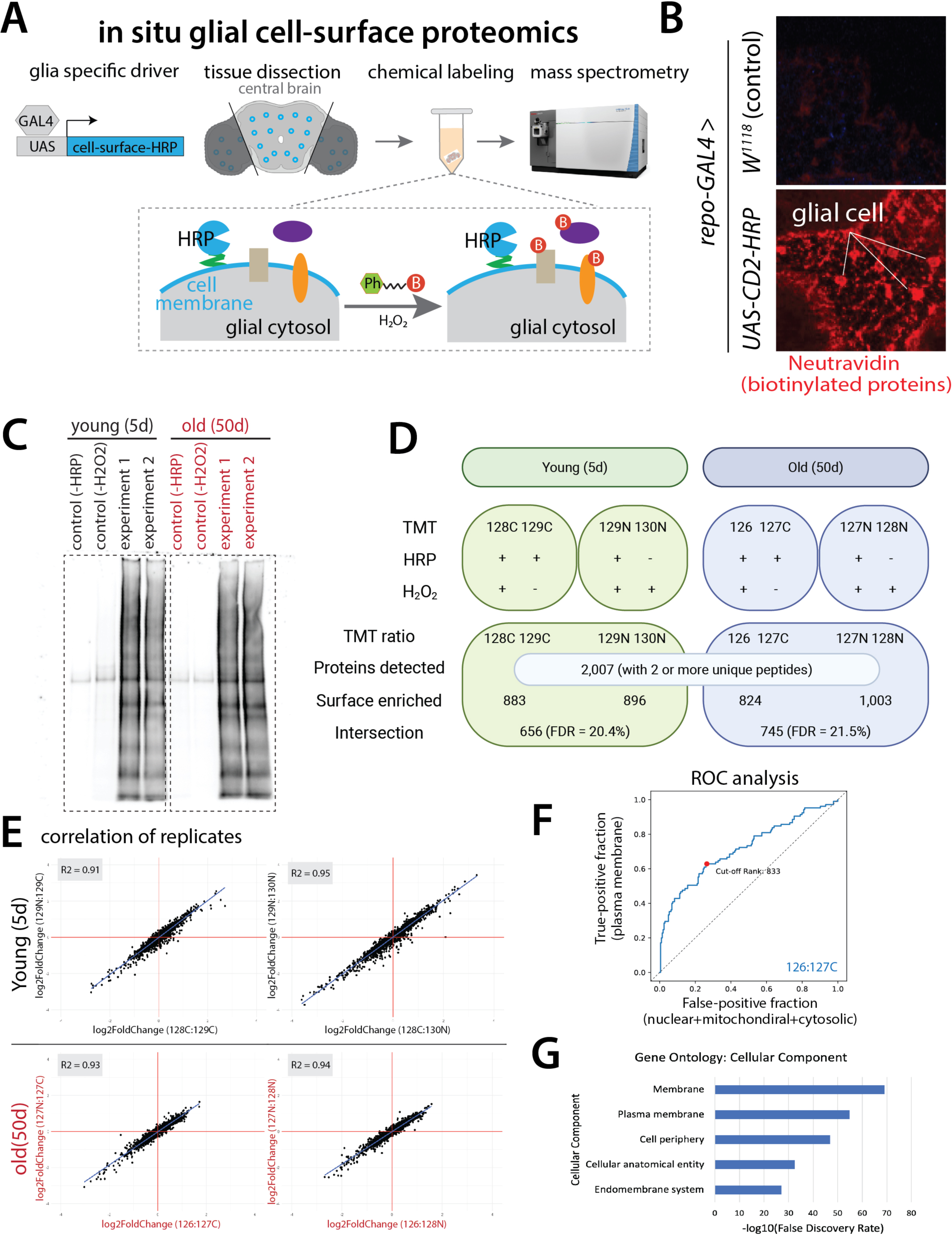
Glial Cell-Surface Proteomics of the Intact Brains of Young and Old Flies. **A:** Transgenic expression of HRP fused to the transmembrane protein CD2 targets CD2-HRP to the extracellular side of the glial plasma membrane. In the presence of H2O2, CD2-HRP converts BxxP substrate into phenoxyl radicals that biotinylate nearby endogenous proteins. Biotinylated proteins are then enriched from brain lysates using streptavidin beads, followed by on-bead trypsin digestion, TMT labeling, and liquid chromatography-tandem mass spectrometry analysis (LC-MS/MS). **B:** To confirm HRP-dependent biotinylation of glial cells, *Repo-GAL4* were crossed with either *UAS-HRP-CD2* or wildtype (w1118) flies and adult brains were stained with neutravidin. Proteins were biotinylated in the experimental group (*UAS-HRP-CD2*) but not in the control group (w1118). **C:** Biochemical characterization by streptavidin blot showed that the glial cell-surface-targeted HRP biotinylated a wide range of proteins compared to the negative control lacking HRP-CD2 expression. **D:** For each time point (5d, 50d), two biological replicates and two negative controls (one lacking HRP-CD2 expression and one lacking H2O2 in the reaction) were profiled, for a total of 8 samples. Each sample consisted of 300 mixed male and female fly brains, for a total of 2,400 brains. In total, we detected 2,007 proteins with two or more unique peptides. Potential contaminants were removed by comparing differential enrichment between the experimental and negative control groups (i.e., the TMT ratio for each protein identified; see Methods). In the TMT labeling system, 126, 127N, 127C, 128N, 128C etc. were used to index and compare different samples. **E:** Strong correlations between biological replicates were observed, further validating the quality of the proteome profiles. **F:** The receiver operating characteristic (ROC) curve depicts the true-positive rate against the false-positive rate of detected proteins, for each biological replicate (only 126:127C pictured). **G:** Gene Ontology (GO) analysis was performed based on the 872 proteins identified following ratiometric and cutoff (see Methods). Top 5 GO terms on cellular compartments with the lowest false discovery rates are shown.

To validate HRP-mediated cell-surface biotinylation of glial cells in intact brains, we crossed *Repo-GAL4* with either *UAS-HRP-CD2* or wildtype (w1118) and stained brains with a fluorophore-conjugated neutravidin, which specifically binds to biotin (Li et al., 2021). We observed extensive HRP-dependent biotinylation of glial cells (**Fig. 1B**). Biochemical characterization by streptavidin blot of the post-enrichment eluate showed that the glial cell-surface-targeted HRP biotinylated a wide range of endogenous proteins compared to the control lacking HRP-CD2 expression (**Fig. 1C**).

### Quantitative comparison of young and old proteomes

We applied this modified cell-surface biotinylation methodology to the glial cells of young (5-day) and old (50-day) *Drosophila* to investigate how glial cell-surface proteomes change as flies age. As the focus of our study was to investigate the more general changes occurring during normal brain aging, we chose to focus on glial cells in the central brain, which is responsible for most of the brain’s higher order functions, including learning and memory, signal integration, behavior, etc. (Robinson et al. 2025). As the optic lobes account for approximately half of all neurons in the adult fly brain and are specialized to process visual stimuli (Robinson et al. 2025), we removed the optic lobes during sample preparation. This was done to prevent biasing findings towards age-related changes in visual function, rather than the more general changes we intended to focus on.

To obtain sufficient cell-surface proteins for downstream mass spectrometry analysis, we dissected 300 mixed male and female fly brains for each sample (for a total of 2,400 brains, see below). To better quantify changes in the glial cell-surface proteome and filter out any contaminants captured in the negative controls, we used an 8-plex tandem mass tag (TMT)-based quantitative strategy (Thompson et al., 2003; Li et al., 2020). We profiled two biological replicates and two negative controls (one lacking HRP-CD2 expression and one lacking H2O2 in the reaction) for each time point (5d and 50d) (**Fig. 1D**). This allowed us to capture endogenously biotinylated proteins, as well as non-specific binders to streptavidin beads. Freshly dissected intact central brains were incubated with the BxxP substrate for one hour before a 5-minute H2O2 reaction. Biotinylated proteins were then enriched from brain lysates using streptavidin beads, followed by on-bead trypsin digestion, TMT labeling, and liquid chromatography-tandem mass spectrometry analysis (LC-MS/MS).

From this 8-plex experiment, we detected a total of 2,007 proteins with 2 or more unique peptides (**Fig. 1D**; Supplementary Table 1) and observed strong correlations between the replicates of each time point (5d and 50d), validating the quality of the proteome profiles (**Fig. 1E**). By comparing differential enrichment between the experimental and negative control groups (i.e., the TMT ratio for each protein), we were able to remove potential contaminants via ratiometric and cutoff analyses (Hung et al., 2014; Li et al., 2020). If a protein had a high TMT ratio, it was considered a true positive (TP), as a high TMT ratio indicates that that protein was extensively biotinylated in the experimental group, but not in the control group. Alternatively, a protein was considered a false positive (FP) if it had a low TMT ratio, as that indicates that protein was similarly biotinylated in both the experimental and the control groups. To calculate the TMT ratios, we paired one biological replicate with one control, as depicted in Fig. 1D.

To determine optimal TMT cutoff values, we plotted the receiver operating characteristic (ROC) curve for each biological replicate (**Fig. 1F**; **Fig. S1B**), which depicts the true-positive rate against the false-positive rate of detected proteins (Li et al., 2020; Nahm, 2022). To maximize the signal-to-noise ratio of the proteomes, each biological replicate was cut off at the position where the value of *true-positive rate (i.e., sensitivity) – false-positive rate (i.e., specificity)* was maximized (described in more detail in the Methods). To further minimize potential contaminants, we selected only TP proteins that overlapped in both biological replicates at each time point (**Fig. 1D**).

In total, we identified 872 proteins following our ratiometric and cutoff analyses, which exhibited high spatial specificity (as seen from the cellular component Gene Ontology (GO) analysis) (**Fig. 1G**). FlyBase, UniProt, and Gene Ontology Data Archive (DOI 10.5281/zenodo.1205166; Version 2023-05-10; Ashburner et al., 2000) were then searched to determine if each of the 872 proteins had plasma membrane and extracellular matrix annotations. We found that of the 872 proteins identified, 682 could be validated by at least one of these databases, further validating the spatial specificity of our cell-surface proteomic labeling (**Fig. S1A**).

### Age-related changes in glial cell-surface proteomes

Of the 872 proteins identified by our glial cell-surface proteomics, we found that 127 were unique to young (5d) flies, 216 were unique to old (50d) flies, and 529 were found in both. This suggests that the cell-surface proteomes of glial cells do in fact shift as flies age; as such, we further compared the glial cell-surface proteomes of young and old flies (**Fig. 2A**).

**Figure 2:**
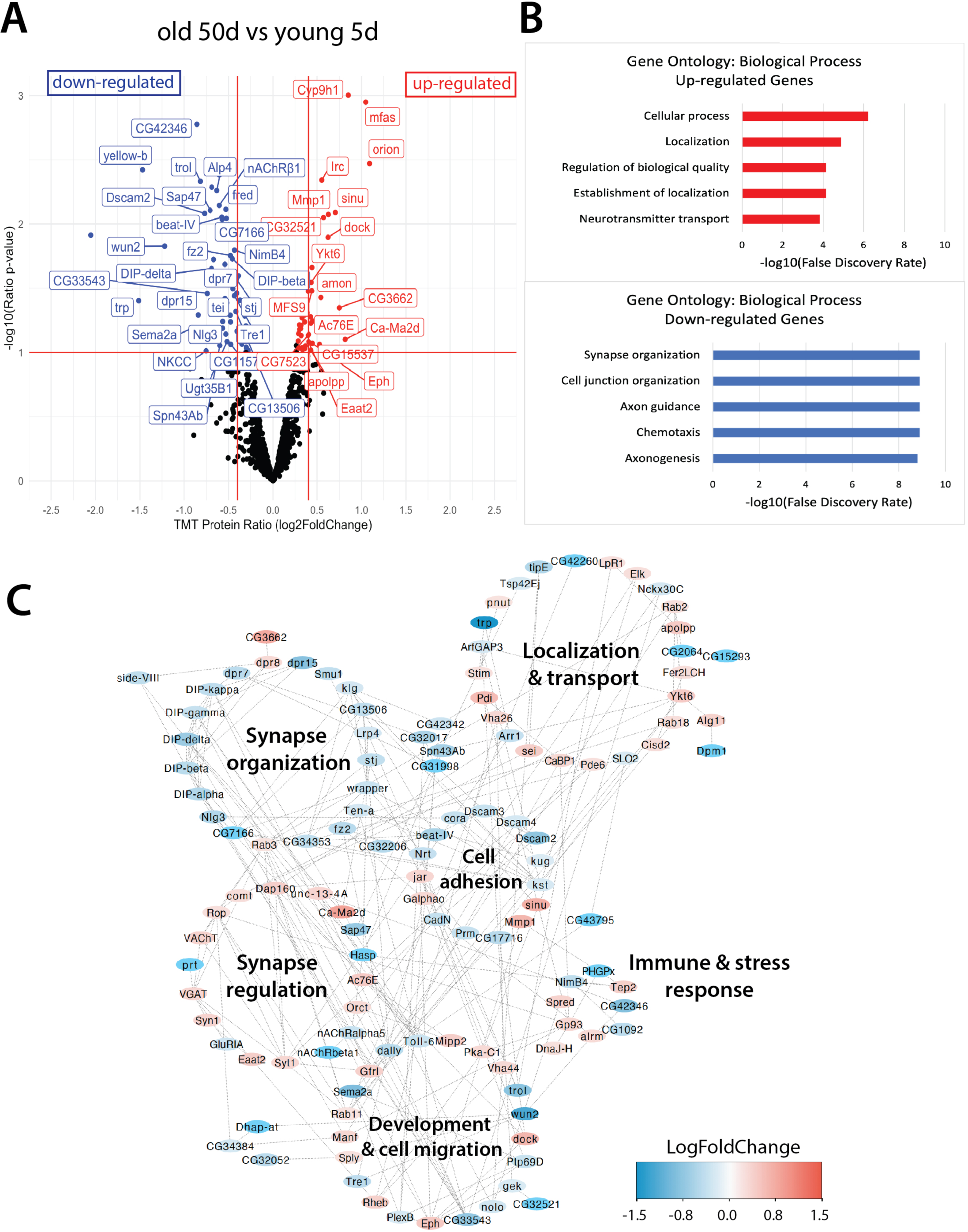
Quantitative comparison of young and old proteomes and selection of candidate genes. **A:** To select approximately 50 candidates for further screening, we generated a volcano plot, where the TMT ratio (log2FoldChange) for each of the 872 proteins identified was plotted on the x-axis and the log10(Ratio P-value) was plotted on the y-axis. Genes associated with the largest change in protein levels between time points were identified using arbitrary cutoff targeting about 50 genes for screening. In total, 48 candidate genes were selected for further screening, most of which have at least one strong human ortholog. Down-regulated candidate genes selected for functional screening are labeled in blue whereas up-regulated candidate genes are labeled in red. **B:** A Gene Ontology (GO) analysis investigating proteome biological processes revealed that genes encoding the most up-regulated proteins were primarily associated with cell localization, transport, and maintaining homeostasis, while most down-regulated proteins were primarily associated with neural development, synapse organization, and neuroplasticity. **C:** Genes encoding the most up- and down-regulated proteins were clustered by their reported protein-protein interactions (PPIs) and corresponding confidence scores (using a Markov clustering algorithm, inflation value set to 2.5). Each protein in the PPI plot was then assessed individually to determine if it had reported function in synapse organization, synapse regulation, cell adhesion, development and cell migration, or localization and transport.

To investigate how old glial cell-surface proteomes change in terms of their biological functions, we performed a GO analysis of genes encoding the most up- and down-regulated proteins. These proteins were selected from the original 872 identified (rather than from the 682 validated by the FlyBase, UniProt, or Gene Ontology Data Archive databases). This was done to ensure we included any potentially important uncharacterized proteins (see Methods). We next plotted the top five retrieved GO terms on biological processes with the lowest false discovery rates (**Fig. 2B**) (Szklarczyk et al., 2023). We found that while most up-regulated genes were associated with cell localization, transport, and homeostasis, most down-regulated genes were associated with neural circuit development, synaptic wiring, and neuroplasticity, suggesting a possible decline in glia-neuron communication as flies age.

Next, we clustered these proteins by their reported protein-protein interactions and corresponding confidence scores using a Markov clustering algorithm (inflation value set at 2.5) (**Fig. 2C**). FlyBase and UniProt were then referenced to determine if each protein in the protein-protein interaction (PPI plot) had previously reported functions in synapse organization, synapse regulation, cell adhesion, development, or localization and transport.

Consistent with the biological functions identified by our GO analyses, we found that PPI clusters for synapse organization, synapse regulation, cell adhesion, and localization/transport were composed primarily of proteins enriched at the same time point, as seen by separately clustered blue or red nodes (**Fig. 2C**). For example, while most nodes in the synapse organization cluster were found to be downregulated in old fly brains, most nodes in the localization and transport cluster were found to be upregulated. Previous studies have found that communities of co-expressed proteins can be linked to disease processes, and that the most strongly correlated proteins or ‘hubs’ within these co-expression modules are often enriched in key drivers of disease pathogenesis (Johnson et al., 2020). As such, we identified the top 10 “hub” genes in our PPI network (**Fig. S1C**). We found that of the top 10 hub genes identified, 7 had reported function in cell adhesion (klg, kst) and synaptic wiring (DIP-β, DIP-δ, DIP-γ, DIP-κ, DIP-ɑ), while the remaining three had reported function in neuron projection (CG33543), axon guidance (CG34353), and dendrite guidance (Toll-6).

To select candidates predicted to be involved in brain aging for functional screening, we identified genes associated with the largest fold-changes in protein levels between time points (5d and 50d) (**Fig. 2A**). Candidate genes were selected for functional screening only if they were known (or predicted) to be located either on the cell-surface or in the extracellular space. In total, we selected 48 candidate genes for functional screening, most of which were known to have at least one strong human ortholog (Supplementary Table 2) (Wang et al., 2017; Wang et al., 2019a; Wang et al. 2019b). Notably, we found that many of these candidates were cell adhesion and neural wiring molecules not previously thought to be particularly active in glia (**Fig. 2B-C**). Interestingly, our final list of candidates included 9 previously uncharacterized proteins (CG genes as Computated Genes), suggesting that future investigations of these proteins may reveal new facets of glial physiology.

### Functional screening of candidate genes predicted to be involved in normal brain aging

To investigate the potential contribution of each candidate gene to normal brain aging, we conducted a broadscale lifespan screen, as previous work has found that brain aging (i.e., age-related neurodegeneration) and lifespan are highly correlated (Mattson et al., 2002; Mattson and Arumugam, 2018). Importantly, several studies have shown that improving brain health can extend lifespan in *Drosophila* (Woodling et al., 2020; Schmid et al., 2022). For our lifespan screen, we used the pan-glial GeneSwitch-GAL4 line initially generated by Nicholson et al. (2008): GSG3285-1, glial-GS hereafter. GeneSwitch-GAL4, a modified, drug-inducible version of the GAL4 system, allows for both spatial and temporal control of transgene expression, which is induced by the drug mifepristone (also known as Ru486) (**Fig. 3A**) (Osterwalder et al., 2001; Nicholson et al., 2008). This allowed us to compare flies from the same genetic background after development was complete, ensuring that our gene manipulations were occurring as flies aged and not during key developmental milestones (Osterwalder et al., 2001). Ultimately, this improved the likelihood that any observed lifespan changes could be attributed to the gene manipulations themselves, rather than unintended developmental defects.

**Figure 3:**
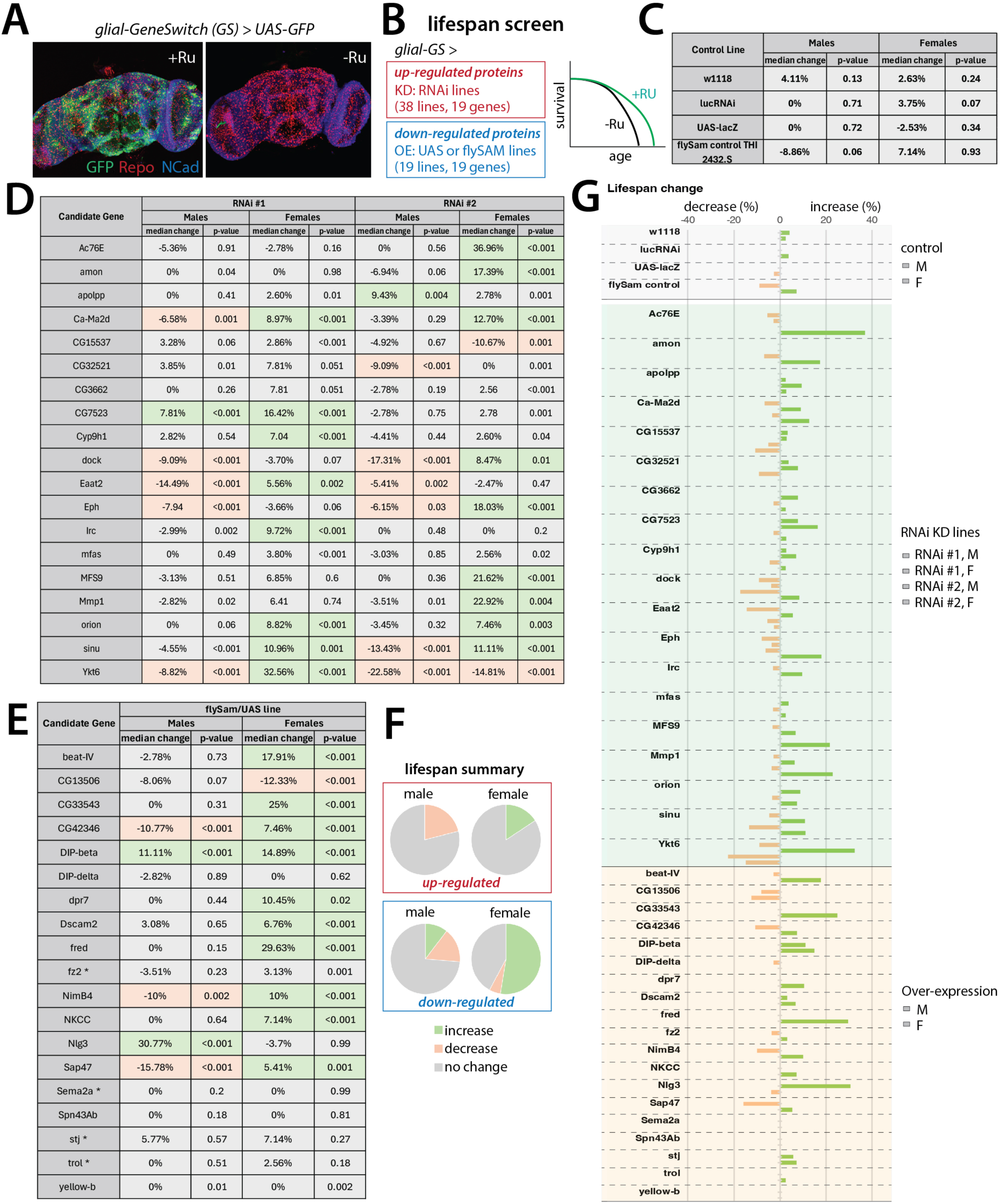
Functional screening of candidate genes predicted to be involved in normal brain aging. **A:** Confocal images showing that the glial-GS line only induced transgene expression in the presence of the inducible drug Ru486. **B:** 48 candidate genes were selected for functional screening. For each of the 19 up-regulated genes, two RNAi lines were used to control for possible off target effects. Of the 29 down-regulated genes selected, we were able to acquire and screen lines for 19 genes. **C:** No obvious lifespan effect was observed in control lines: a w1118 wildtype control, a UAS-luciferase RNAi control, a UAS-lacZ control, and a flySAM control. **D:** Pie charts summarizing the results of our broadscale lifespan screen for male and female flies. **E:** Of the 19 up-regulated candidate genes screened, RNAi knockdown of 4 genes led to consistent lifespan decreases for males. Alternatively, RNAi knockdown of 3 genes led to consistent lifespan increases for females. **F:** Of the 19 down-regulated candidate genes screened, overexpression of 2 genes led to lifespan increases for males, whereas overexpression of 3 genes led to lifespan decreases. In parallel, overexpression of 10 genes led to lifespan increases for females, whereas overexpression of 1 gene led to decreased lifespan.

For each of the 19 up-regulated candidate genes, we screened 2 RNAi transgenic lines to control for potential off-target effects (**Fig. 3B**; RNAi lines used for each gene are detailed in Supplementary Table 3) (Dietzl et al., 2007; Ni et al., 2011; Perkins et al., 2015). For each down-regulated candidate gene, we screened one over-expression transgenic line (**Fig. 3B**; Supplementary Table 3), using either UAS-cDNA lines or flySAM2.0 lines. flySAM is a CRISPRa (CRISPR activation) system used in *Drosophila* to activate gene expression (Jia et al., 2018). flySAM2.0, an updated and simplified version of flySAM1.0, increases experimental efficiency by combining UAS-dCas9 and sgRNA in the same transgenic fly, so that any flySAM2.0 lines can be directly crossed with a GAL4 line (Jia et al., 2018). As flySAM lines have been found to recapitulate overexpression phenotypes comparable in strength to UAS-cDNA lines, they present an ideal alternative to investigate candidates for which no UAS-cDNA line is currently available (Jia et al., 2018). For the down-regulated genes, we were able to acquire and screen lines for 19 of the 29 candidate genes (**Fig. 3B**).

First, we confirmed that the glial-GS line only induced transgene expression in the presence of the inducible drug Ru486. We found that the glial-GS drove *UAS-CD8-GFP* expression in the presence of the inducible drug Ru486, but not in the presence of the control (**Fig. 3A**), consistent with the previous study (Nicholson et al., 2008). Next, we crossed the glial-GS line with several control lines (a w1118 wildtype control, a UAS-luciferase RNAi control, a UAS-lacZ control, and a flySAM control). Importantly, we did not find an effect of the Ru486 drug itself on control lifespan for this glial-GS line (GSG3285-1) (**Fig. 3C**; **Fig. 4C-D; Fig. S2A-B**).

**Figure 4:**
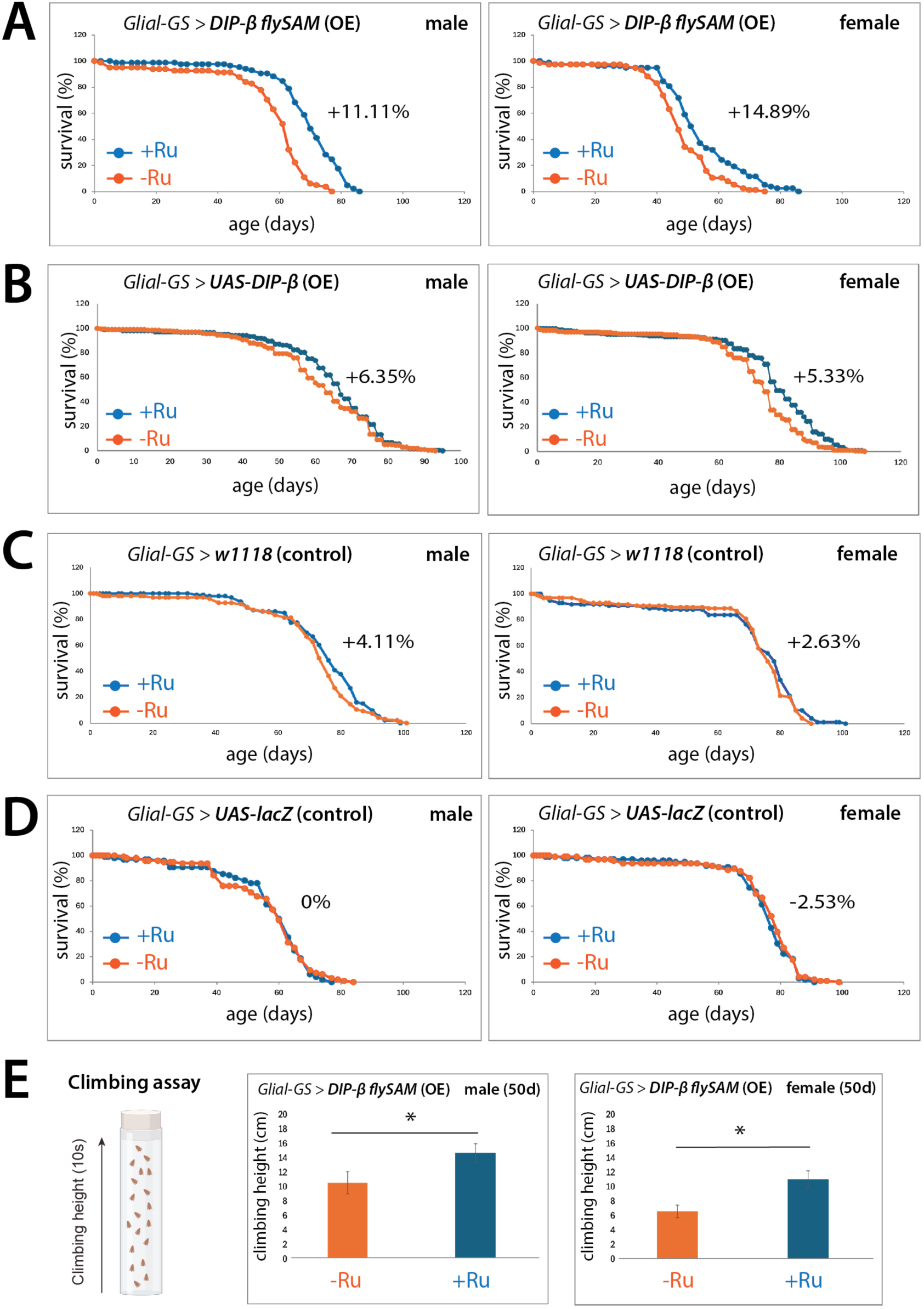
Validating that pan- glial overexpression of DIP-β increases lifespan and serves a protective function during aging. **A-D:** Glial overexpression of DIP-β (using both a flySAM2.0 and a UAS overexpression line) significantly extends lifespan for both male and female flies, compared to the negative controls (only the w1118 and UAS-lacZ controls are shown here). **E:** To further validate that glial overexpression of DIP-β is serving a protective function during brain aging, climbing (i.e., negative geotaxis) assays were conducted. Glial overexpression of DIP-β was also associated with improved climbing and motor control ability in aged flies.

We observed sex-specific lifespan effects for many of these screened genes. Of the 19 up-regulated genes that were screened, we found that RNAi knockdown of 3 genes (*Ca-Ma2d, orion, sinu*) led to consistent increases in lifespan for females but not for males (**Fig. 3D-E; Fig. S3-6**). We also found that RNAi knockdown of 4 genes (*dock, Eaat2, Eph, Ykt6*) led to consistent decreases in lifespan for males (i.e., lifespan decreases were observed for both RNAi lines used), but inconsistent lifespan changes were observed for females (**Fig. 3D-E; Fig. S3-6**). Interestingly, we did not find any RNAi lines that increased lifespan for both males and females. Of the 19 down-regulated genes that were screened, we found that overexpression of *Nlg3* led to increases in lifespan for males, but not for females, whereas overexpression of 3 genes (*CG42346, NimB4, and Sap47*) led to lifespan decreases for males, but lifespan increases for females (**Fig. 3D, 3F; Fig. S7-8**). Alternatively, we found that overexpression of 6 genes (*beat-IV, CG33543, dpr7, Dscam2, fred*, and *NKCC*) led to increases in lifespan for females but observed no changes in lifespan for males (**Fig. 3D, 3F; Fig. S7-8**). These sex-specific differences in lifespan is an interesting subject for future validation and investigation. Next, we focused on DIP-β, which was the only candidate gene associated with consistent increases in lifespan for both male and female flies (**Fig. 4A-B**).

### Validating that glial overexpression of DIP-β increases lifespan

Following our functional screen, we identified DIP-β as the most promising candidate for further investigation. Interestingly, DIP-β was also identified as the top-ranked “hub” gene in our PPI analysis of the most up- and down-regulated genes (**Fig. S1C**).

To validate these initial findings, we conducted two additional lifespan trials, again using the DIP-β flySAM2.0 line to overexpress DIP-β in glia. We observed consistent increases in lifespan across DIP-β flySAM2.0 lifespan trials (**Fig. 4A**). Next, we conducted two more lifespan trials, this time using a UAS-DIP-β line to overexpress DIP-β in glia. Again, we observed consistent increases in lifespan across UAS-DIP-β lifespan trials (**Fig. 4B**). All control lines did not show a significant lifespan effect (Fig. 4C, D), indicating that the lifespan increases observed when DIP-β is overexpressed in glia are likely due to the DIP-β expression manipulations themselves, rather than the Ru486 inducer. While the lifespan extensions observed for glial *DIP-β* overexpression were relatively modest, their reproducibility across different overexpression platforms suggests they are biologically relevant, even if they do not reach the dramatic shifts seen in some caloric restriction or drug-based models (Miquel et al., 1976; Partridge et al., 2005; Burger et al., 2006).

As we observed greater lifespan extensions for DIP-β flySAM2.0 than UAS-DIP-β, we chose to use this line for our subsequent investigations. This decision was further bolstered by prior studies, which indicate that flySAM lines target all the isoforms of a given protein, whereas UAS-cDNA lines are specific to only one isoform (Zirin et al., 2020). Together, this suggests that DIP-β flySAM2.0 is likely the more efficient overexpression line, as DIP-β has several known isoforms (FlyBase). While we did observe a shorter lifespan for the DIP-β flySAM2.0 negative (i.e., ethanol) controls, compared to the UAS-DIP-β negative controls, we found that our flySAM2.0 lines exhibited shorter baseline lifespans overall (compared to the traditional UAS-cDNA lines; **Fig. S7-8**). The shorter baseline lifespans we observed for our flySAM2.0 lines are likely due to the specific genetic background of the flySAM transgenic insertions, as low levels of “leaky” expression have been previously reported for these lines (Jia et al., 2018). However, we believe that the lifespan extensions we observed for DIP-β flySAM2.0 is a robust biological effect, rather than an artifact of reduced viability for the following reasons. First, by utilizing the GeneSwitch (GS) system, we were able to compare the lifespan of flies with the exact same genetic background (+/- RU-486). This ensures that the extension we report is specifically due to the induction of the transgene itself, rather than a comparison between disparate lines with different basal fitness levels. Second, if the lifespan extensions were merely a recovery from lower baseline viability, we would expect to see similar improvements across other flySAM2.0 lines in our screen. However, DIP-β was the only candidate across our screen that significantly increased lifespan in both sexes (Extended Data Figs. 7 & 8). Third, the lifespan-extending effect of DIP-β was independently confirmed using a traditional UAS-cDNA line, which importantly does not share the same baseline viability issues as the flySAM2.0 lines.

Next, we conducted climbing (i.e., negative geotaxis) assays, as motor control and climbing ability are commonly considered reliable phenotypic indicators of brain aging and age-related neurodegeneration (Barone and Bohmann, 2013; Beharry et al., 2013; Manjila and Hasan, 2018). We found that pan-glial overexpression of DIP-β was also associated with improved climbing and motor control ability in aged flies for both males and females (**Fig. 4E**). Altogether, these results indicate that DIP-β can play an important role in maintaining healthy brain function and delaying phenotypes associated with age-related neurodegeneration.

### Investigating the downstream transcriptomic effects of overexpression of DIP-β in glial cells during aging

To investigate the downstream transcriptomic effects of upregulating glial expression of DIP-β as flies age, we conducted single-nucleus RNA sequencing (snRNA-seq). To do this, we prepared two whole-head samples collected from old (50d) flies: one where DIP-β had been overexpressed in glia from day 5-50 and one where DIP-β expression had not been manipulated (**Fig. 5A**).

**Figure 5:**
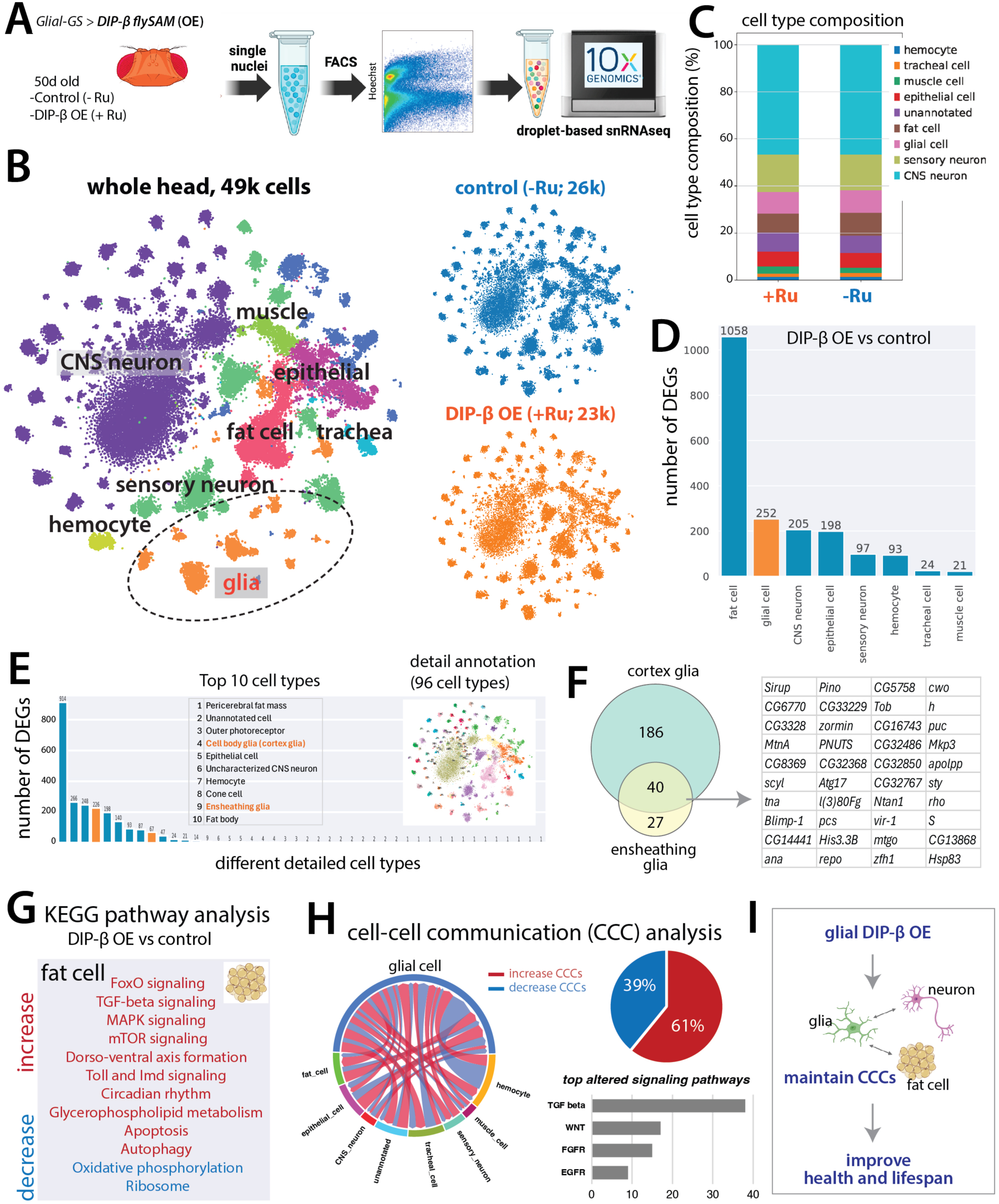
Single-nucleus RNA-sequencing - investigating the downstream transcriptomic effects of glial overexpression of DIP-β during aging. **A:** Two whole-head samples (20 heads each) were prepared for snRNA-seq: one where DIP-β had been overexpressed in glia from day 5-50 and one control. Single-nucleus RNA-seq libraries were prepared using the 10X Genomics Chromium Single Cell 3’ HT v3.1 protocol. **B:** In total, approximately 49,000 high-quality cells were obtained after quality control and filtering from our fly head samples, which consist of 8 broad cell types. **C:** Broad cell-type cluster compositions were largely consistent between the glial DIP-β overexpression treatment (Ru486+) and the control (Ru486-) groups. **D:** A broad analysis of differentially expressed genes (DEGs) revealed that the glial cell-type cluster showed the second highest number of DEGs between the glial DIP-β overexpression treatment (Ru486+) and the negative control (Ru486-) groups. **E:** A more detailed analysis investigating DEGs across 90 different cell subtypes revealed that of the glial cell subtypes, cortex glia (i.e., “adult brain cell body glia”) and ensheathing glia showed the highest number of DEGs between the DIP-β overexpression treatment (Ru486+) and the control (Ru486-) groups. **F:** Of the 252 DEGs identified for glial cells, 186 were unique to cortex glia, 27 were unique to ensheathing glia, and 40 were found to be differentially expressed in both cortex and ensheathing glia. **G:** As fat cells exhibited the highest number of DEGs between the DIP-β overexpression and control groups, we conducted a KEGG analysis to evaluate the biological functions of these genes. Most fat cell DEGs were up-regulated, and the top altered metabolic and signaling pathways for these genes were found to be related to homeostasis, signal integration, stress resistance, and longevity. **H:** Cell-cell communication (CCC) analysis was performed using FlyPhoneDB2. DIP-β overexpression was generally associated with an increase in cell-cell communication, particularly glia-neuron and glia-fat cell communication. The top altered glial signaling pathways are shown. **I:** Our proposed model: Glial overexpression of DIP-β leads to improved glia-neuron and glia-fat cell communication as flies age, which may contribute to the observed lifespan extensions.

For our snRNA-seq, we used the DIP-β flySAM2.0 to overexpress DIP-β in glia (as opposed to the UAS-DIP-β line), as we consistently observed stronger lifespan-extension effects for the flySAM2.0 line. Additionally, we chose to sequence the heads of female flies, as we consistently observed stronger lifespan-extension effects for females. Sequencing whole-head samples allows for detailed profiling of glia, neurons, and other related cell types and subtypes (including fat cells, muscle, trachea, epithelial cells, etc.). As such, this enables us to form a more comprehensive understanding of global changes across the entire fly head (as previously demonstrated in our Aging Fly Cell Atlas (Lu et al., 2023) and Alzheimer’s Disease Fly Cell Atlas (Park et al., 2025).

In total, we obtained 48,959 high-quality cells after the quality control steps: ∼23,000 from the DIP-β overexpression (i.e., Ru486+) sample and ∼26,000 from the control (i.e., Ru486-) sample (**Fig. 5B**). Based on our Fly Cell Atlas cell annotations (Li et al., 2022), these cells were annotated into 8 broad cell types (CNS neurons, sensory neurons, glia, fat cells, hemocytes, muscle, trachea, and epithelial cells) and 96 detailed cell subtypes (**Fig. S9**).

First, we compared broad cell type compositions between the DIP-β overexpression and the control samples. We found that broad cell type compositions were largely consistent between the two groups (**Fig. 5C**), suggesting that DIP-β overexpression does not change cell ratios within the head. Next, we analyzed differentially expressed genes (DEGs) for each of the 8 broad cell types. We found that glial cells exhibited the second highest number of DEGs after fat cells (**Fig. 5D**). Further analysis of DEGs across 96 detailed cell subtypes revealed that cortex glia (i.e., “adult brain cell body glia”) and ensheathing glia were among the top affected cell subtypes (**Fig. 5E**). Between these two glial subtypes, 186 DEGs were unique to cortex glia, 27 were unique to ensheathing glia, and 40 were found in both cortex glia and ensheathing glia (**Fig. 5F**).

As fat cells exhibited the highest number of DEGs between the glial DIP-β overexpression and control samples, we conducted a KEGG pathway analysis to evaluate the biological functions of these genes (**Fig. 5G**) (Chen et al., 2017). We found that fat cells showed more up-regulated DEGs than down-regulated ones, and that the top altered metabolic and signaling pathways for these up-regulated genes were related to homeostasis, signal integration, stress resistance, and longevity (**Fig. 5G**). Interestingly, these results are consistent with our previous findings that fat cells are often among the cell types found to be most sensitive to changes related to aging and neurodegeneration (Lu et al., 2023; Park et al., 2025).

Next, we conducted a cell-cell communication (CCC) analysis to evaluate how glial overexpression of DIP-β impacts cell-cell signaling pathways during normal brain aging (**Fig. 5H**) (Wilk et al., 2024). To do this, we employed the FlyPhoneDB2 that we recently developed (Qadiri et al., 2025). FlyPhoneDB2 is a computational platform for estimating cell-cell signaling using *Drosophila* snRNA-seq data and a curated list of high-confidence ligand-receptor pairs (Qadiri et al., 2025; Liu et al., 2021). By aggregating and comparing ligand-receptor expression values for various groups of cells, we were able to infer which groups of cells are likely interacting with glia and which ligand-receptor pairs are contributing to the glial-other cell communication changes. We found that DIP-β overexpression was generally associated with an increase in cell-cell communication compared with the aged control, particularly glia-neuron and glia-fat cell communication (**Fig. 5H**). More specifically, we found that DIP-β overexpression led to increases in expression for 61% and decreases in expression for 39% of ligands and receptors. Additionally, we found that the top altered glial signaling pathways include TGF-beta, WNT, FGFR, and EGFR signaling pathways (**Fig. 5H**).

Combined, these results suggest that glial overexpression of DIP-β leads to improved glia-neuron and glia-fat cell communication as flies age, which may be contributing to the observed lifespan extensions through multiple signaling pathways (**Fig. 5I**). As such, it will be interesting to investigate the molecular mechanisms underlying how DIP-β overexpression leads to these signaling changes, how DIP-β’s binding partners (multiple cell-surface Dpr proteins, see Discussion) are involved, and which downstream pathways play key roles in mediating the lifespan extension in future studies.

## Discussion

Age-related changes in the cell-surface proteomes of glial cells are hypothesized to play important roles in the dysregulation of glia-neuron interactions observed in brain aging (Barres, 2008; Kremer et al., 2017). However, these changes have not been previously characterized due to technical limitations. Here, we adapted a platform to profile spatiotemporally resolved glial cell-surface proteomes from intact brains in young and old *Drosophila* (Li et al., 2020). Our study revealed global downregulation of synaptic wiring and cell adhesion molecules in the aged brain, and identified several molecules not previously thought to be particularly active in glia. Additionally, our study identified the neural wiring molecule DIP-β as a potential regulator of healthy brain function during brain aging. Here, we explore how cell adhesion and synapse organization molecules like DIP-β may be playing a role in age-related neurodegeneration.

In *Drosophila*, several superfamilies of cell adhesion molecules (CAMs) are known to contribute to neural circuit organization (including cadherins, immunoglobulins, leucine-rich repeat proteins, and teneurins) (Byron et al., 2010; Bulgakova et al., 2012; Sanes and Zipursky, 2020; Xu et al., 2024). While the mechanisms underlying precise neural wiring are not yet fully understood, it is generally thought that each subset of neurons expresses a unique combination of CAMs, which act as highly specified identification tags (Elmariah et al., 2005; Lin and Koleske, 2011; Silies and Klambt, 2011; Carrillo et al., 2015; Tan et al., 2015; Cosmanescu et al., 2018; Li et al., 2020; Kim et al., 2020). Indeed, previous work has provided ample evidence in support of this, demonstrating that neuronal cell subtypes often express the same groups of CAMs, and that loss-of-function mutations in these molecules result in profound (and often predictable) neural wiring defects (Carrillo et al., 2015; Wang et al., 2022; Ma et al., 2023). While many previous studies have painstakingly delineated the contributions of various CAMs to precise synaptic wiring in the developing brain, these molecules do not have a comparably well-defined role in the aging brain (Lin and Koleske, 2010; Ma et al., 2023).

Global downregulation of synaptic wiring molecules, loss of cell identity, and diminished synaptic connectivity are all considered hallmarks of age-related neurodegeneration (in both normal brain aging, as well as in a wide variety of neurodegenerative diseases) (Lin and Koleske 2010; Li et al., 2020; Banerjee et al., 2021; Banerjee et al., 2024; Li et al., 2022). While most of what glia do in the brain is thought to depend on their close physical relationship with neurons, synapses, and other glial cells, we know surprisingly little about how glia precisely associate and coordinate with their intended targets (Silies and Klambt, 2011; Freeman, 2015; Cho et al., 2018; Bittern et al., 2021). For example, exactly how one cortex glial cell is able to accommodate the unique needs of up to 100 neurons simultaneously remains one of the greatest mysteries in glial cell biology (Hillen et al., 2018; Chen et al., 2024). Previous studies have demonstrated that at least some CAMs play important roles in glia-neuron interactions (Grumet and Edelman, 1988; Lepeta et al., 2016; Chou et al., 2020); for example, Neurexins and Neuroligins are known to help coordinate astrocyte-synapse contact sites during synaptogenesis (Grumet and Edelman, 1988; Lepeta et al., 2016; Chou et al., 2020; Chen et al., 2024). Additionally, CAMs have been found to function in a range of other biological processes related to trans-synaptic signaling, including synaptic vesicle organization, receptor clustering, and structural and functional plasticity (Chou et al., 2020). However, we do not yet fully understand the role these CAMs may be playing in enabling glia to effectively interact with, support, and coordinate with other cells in the brain.

In *Drosophila*, heterophilic binding between members of two immunoglobulin sub-families, the 11-member DIP (Dpr interacting proteins) family and the 21-member Dpr (defective proboscis extension response) family, are known to regulate precise synapse organization (Carrillo et al., 2015; Tan et al., 2015; Cosmanescu et al., 2018; Xu et al., 2019; Honig and Shapiro, 2020; Sanes and Zipursky, 2020; Sergeeva et al., 2020; Wang et al., 2022; Ma et al., 2023). While DIPs and Dprs are best known for regulating precise synaptic wiring in the developing optic lobe, some studies have found that DIP/Dpr proteins also play important roles in the central adult brain (Carrillo et al., 2015; Wang et al., 2022; Ma et al., 2023). For example, one study found that like optic lobe neurons, dopaminergic and clock neurons express cell-subtype-specific combinations of DIP and Dpr proteins, and that cyclic expression of DIP-β in a small subset of clock neurons (small LNvs) plays an important role in sleep onset and duration in adult flies (Ma et al., 2023). As transcript cycling across the day has been well documented in clock neurons, it is possible that DIP-β may be playing a role in circadian-related changes in synaptic plasticity (Ahmad et al., 2021). In support of this, the human ortholog of DIP-β, IgLON5, is involved in processes related to neuroplasticity and neurogenesis, and is additionally suspected of playing a role in maintaining blood-brain barrier (BBB) integrity (Madetko et al., 2022; Zhang et al., 2023). Recently, IgLON5 was implicated in a rare autoimmune disease, anti-IgLON5, a condition that presents with several nonspecific symptoms, including sleep disturbances, gait abnormalities, and impaired cognitive function (Madetko et al., 2022; Zhang et al., 2023). Combined, these findings suggest that DIP-β may be playing important roles in neuroplasticity, sleep regulation, and BBB permeability.

Interestingly, we found that of the 32 DIP and Dpr proteins known in *Drosophila*, nine (DIP-β, DIP-δ, DIP-γ, DIP-κ, DIP-ɑ, Dpr7, Dpr 8, Dpr12, Dpr 15) were included in our final cell-surface proteomes. Of these 9 proteins, four (DIP-β, DIP-δ, Dpr8, Dpr15) were additionally included in the 48 candidates selected for further screening, suggesting that these molecules likely play important roles in age-related neurodegeneration. Previous work has demonstrated that, in general, DIPs bind preferentially to Dprs (Sergeeva et al., 2020), and that while neurons typically express several Dprs, DIPs are expressed far more selectively (1-2 per neuron) (Cosmanescu et al., 2018; Wang et al., 2022). DIPs and Dprs have been found to display a broad range of binding affinities, as some bind heterophilically to one partner, others are more promiscuous, and a few bind homophilically (Cosmanescu et al., 2018; Sanes and Zipursky, 2020; Xu et al., 2019). Of the 11 DIP proteins, DIP-β has been found to display some of the strongest and most promiscuous binding affinities (i.e., DIP-β binds strongly to a broader range of Dprs, such as Dpr6, Dpr8, Dpr9, Dpr15, and Dpr21, than most other DIPs) (Cosmanescu et al., 2018; Sergeeva et al., 2020). This suggests that DIP-β may have the potential to interact with a broader range of neuronal cell types than other DIP proteins. While another down-regulated DIP (DIP-δ) was also included in our functional screen, we did not observe any lifespan increases when DIP-δ was overexpressed in glia. This could be because DIP-δ reportedly binds with only one Dpr (Dpr 12) and thus may be interacting with a narrower range of neurons than DIP-β (Honig and Shapiro, 2020).

As one of the best documented roles of glial cells is to facilitate controlled transport of nutrients and macromolecules, both into and out of individual neurons via the blood-brain barrier (BBB) (Artiushin et al., 2018; Moulton et al., 2021; Axelrod et al., 2023; Contreras and Klambt, 2023), our finding that upregulation of DIP-β in glia primarily impacts cortex and ensheathing glia is particularly interesting. While cortex glia are thought to facilitate the efficient transfer of nutrients from the hemolymph to neuronal cell bodies, ensheathing glia clear debris from the neuropil (Hartenstein, 2011; Freeman, 2015). As previous work suggests that DIPs and Dprs are primarily expressed in neurons under normal conditions, we suspect that glial overexpression of DIP-β, a CAM with known promiscuous binding affinities, may have partially compensated for the global downregulation of other glial CAMS in the aging fly brain. This could help explain why glial DIP-β overexpression had such a broad impact on the transcriptomes of so many different cell types. Altogether, our findings suggest that overexpressing DIP-β in glia may help preserve the ability of these cells to interact with and support their intended targets in the aging brain. However, we also acknowledge the possibility that the differential gene expression between control and DIP-β overexpression flies may reflect transcriptional changes, rather than altered glia-neuron communication *per se*.

Interestingly, we observed sex-specific lifespan effects for many of our candidate genes, as female flies were far more likely than males to show increases in lifespan following our gene expression manipulations (Fig. 3). Previous work has found that the GeneSwitch inducer, Ru486, can have sex-specific effects on metabolism and lifespan, depending on the nutritional environment (Dos Santos & Cocheme, 2024). Specifically, Ru486 has been shown to counteract the lifespan-shortening effects of mating in females, an effect less pronounced in males (Landis et al., 2015; Tower et al., 2017). While we optimized our media (i.e., nutritional environment) and used the *GSG3285-1* line to minimize these potential baseline effects, it remains possible that certain genotypes exhibited a sex-specific sensitivity to the inducer itself. However, it is important to note that Ru486 did not extend mated female lifespan for any of our control lines.

Beyond technical considerations of our *GSG3285-1* line, sex differences in aging are well-documented in *Drosophila* and other organisms (Regan et al., 2016; Austad & Fischer, 2016). Male and female flies exhibit distinct transcriptional trajectories and metabolic shifts as they age. Furthermore, recent studies have highlighted that glial function and the neuroinflammatory landscape can differ significantly between sexes, which may dictate how a specific genetic manipulation impacts the aging process in a sex-dependent manner (Müller et al., 2025). While our screen identifies *DIP-β* as a rare candidate that extends lifespan in both sexes, the prevalence of female-specific hits in our data suggests that the female “aging program” may be more plastic or responsive to the specific glial pathways we targeted. These observations provide a valuable foundation for future studies into the mechanisms of sex-specific neuroprotection.

Importantly, our selection of the specific GeneSwitch line we used for this study was based on several critical experimental considerations: 1.) To minimize background toxicity. We initially tested multiple *Repo-GeneSwitch* lines; however, we found these lines exhibited significant, genotype-dependent lifespan reductions upon Ru486 administration, even in control crosses. This baseline toxicity confounded the interpretation of any potential lifespan effects. *GSG3285-1* was chosen for this study, as it provided a robust control baseline and didn’t show lifespan effects with Ru486 treatment in multiple control lines. This is essential for lifespan studies. 2.) The driver breadth and specificity. As noted in its original characterization (Nicholson et al., 2008) and a later study (Catterson et al. 2023), *GSG3285-1* is a powerful tool for pan-glial induction, though it may include a small population of sensory neurons. Furthermore, while *Repo* is a standard glial marker, it does not label all glial subtypes with equal intensity according to our experience. As such, the “non-overlapping” signal observed in Fig. 3A may reflect this staining bias. 3.) The expression mosaicism. The fact that some glial cells do not show GFP expression in Fig. 3A suggests a degree of mosaicism, which is common to many GeneSwitch lines (Osterwalder et al., 2001). While we acknowledge this means our manipulations may target a broader subset — rather than every single glial cell — the fact that we still observed significant lifespan effects across two independent platforms (UAS and CRISPRa) suggests that the targeted population is sufficient to mediate these systemic effects.

In summary, our study revealed that the maintenance of cell-cell communication and healthy brain function during aging relies on several previously unexpected cell-adhesion and synaptic wiring molecules. Our findings highlight the power of using *in-situ* cell-surface proteomics to capture the native complexity of proteomes that would otherwise be lost using more traditional proteomics methods. Further investigations of these CAMs (particularly DIP-β) will be instrumental for expanding our understanding of the mechanisms underlying how these molecules contribute to synaptic connectivity and the maintenance of healthy brain function during normal brain aging.

## ACKNOWLEDGMENTS

We thank the Li lab members for their constructive advice on this project. We thank Dr. Chundi Xu for sharing the UAS-DIP-Beta fly line.

## AUTHOR CONTRIBUTIONS

Conceptualization: M.M., H.L.

Fly experiments: M.M, E.H., O.P., A.V.

Cell-surface proteomics: J.L., H.L., L.L., N.D.U., D.K.C., D.R.M, S.A.C

Data analysis: M.M., B.S., M.C.W., T-C.L., M.Q., Y.H.

snRNA-seq: T.J., Y-J.P., Y.Q.

Transgenic fly: J.Z., N. P.

Writing: M.M., H.L.

Review and Editing: All authors. Resource: L.L., K.V., H.L.

Supervision: H.L.

Funding Acquisition: H.L.

## Methods

### Lead contact and materials availability

Further information and requests for resources and reagents should be directed to the Lead Contact, Hongjie Li. All unique reagents generated in this study are available from the Lead Contact.

### *Drosophila* as a Model System

*Drosophila* have long been established as one of the best developed model species for studying nervous system biology (Freeman, 2015; Jourjine and Hoekstra, 2021). This is largely due to their short generation time, and to their powerful and extensive molecular genetics toolkit, which enables investigators to single out, manipulate, and quantitatively compare genetically and morphologically defined cell populations (Jourjine and Hoekstra, 2021). Ultimately, this allows investigators to carry out broadscale screens of many cells and genes rapidly and affordably, after which the findings can be translated into mouse and human models. Additionally, *Drosophila* have proven a prime model system for studying aging and age-related diseases, as many key aging pathways are known to be conserved from flies to mammals (Giansanti and Piergentili, 2025; Park et al., 2025).

In *Drosophila*, glial cells follow neurons as the most abundant cell type in the brain and are thought to represent approximately 5-10% of the total population of cells within the fly central nervous system (CNS) (Freeman, 2015). A large body of work has found evidence that *Drosophila* glia share many morphological and functional similarities with mammalian glia, and that this conservation of basic glial cell biology extends to the molecular level (Freeman, 2015). In *Drosophila*, the CNS is composed primarily of three glial cell subtypes (cortex glia, ensheathing glia, and astrocyte-like glia), which fully envelope the CNS scaffold of neuron cell bodies, neurites, and synapses (Freeman, 2015). Recent studies have found that these CNS glial cells play several key roles in the *Drosophila* brain, including homeostasis, plasticity, neurodegeneration, etc. (Freeman, 2015).

### Drosophila stocks and genotypes

All fly stocks were kept in a humidified, temperature controlled incubator at 25℃ with a 12:12 hour light/dark cycle (Bauer et al., 2004; Li et al., 2020). Fly stocks were housed in vials containing standard cornmeal medium and flipped to new food vials every 2-3 weeks. Standard cornmeal medium consisted of cornmeal, agar, inactive dry yeast, molasses (Genesee Scientific), and water, as well as nipagin (dissolved into 140-proof ethanol, Decon Labs) and propionic acid (Sigma Aldrich). In cases where larger numbers of flies of a certain genotype were required for experiments (such as our glial-GS line), flies were transferred to bottles containing standard cornmeal medium and flipped to new food bottles every 2-3 weeks. The following lines were used: w1118, UAS-luciferase RNAi, UAS-lacZ, Repo-GAL4, UAS-redStinger, UAS-CD4-GFP (Han et al., 2011), UAS-CD8-GFP (Lee and Luo, 1999), UAS-HRP-CD2 (Larsen et al., 2003), GSG3285-1 GeneSwitch (Nicholson et al., 2008), UAS-DIP-β (Xu et al., 2019). All UAS and RNAi lines were generated previously (Xu et al., 2019) and acquired from the Bloomington *Drosophila* Stock Center and the Vienna *Drosophila* Resource Center (Supplementary Table 3). All flySAM2.0 overexpression and flySAM control lines were generated by the Perrimon Lab (Jia et al., 2018).

### Glial cell-surface biotinylation in intact brains of young and old flies

Glial cell-surface proteins were labeled in intact male and female fly brains, using the biotinylation methods previously described by Li et al. 2020 (Cell). Brain dissections were performed using Dumont #5 Forceps (Fine Science Tools). All dissection tools were cleaned prior to brain dissections with Milli-Q ultrapure water (EMD Millipore) to remove any detergent or chemical contaminants. Brains were then dissected in Schneider’s medium (Thermo Fisher), which had been pre-cooled on ice, and subsequently transferred into 1.5 mL protein low-binding tubes (LoBind, Eppendorf), which contained 500 μL of Schneider’s medium per sample (i.e., EP tube) and were kept on ice. Optical lobes were removed, and only central brains were used. Following dissection, brains were briefly rinsed with fresh Schneider’s medium to remove fat bodies and dissection debris. Next, 100 μM BxxP was dissolved in fresh Schneider’s media by extensive vortex and sonication. Brains were then incubated in the BxxP-Schneider’s medium for 1 hour on ice (i.e., pre-loaded with BxxP), with occasional mixing by pipetting (Durojaye, 2021). Following incubation, 1 mM (0.003%) H2O2 (Thermo Fisher) was added to the BxxP-Schneider’s medium for 5-minutes at room temperature, allowing H2O2 to form a reactive complex with HRP (HRP-H2O2) and oxidize BxxP into phenoxyl radicals that react with endogenous proteins in very close proximity (Durojaye, 2021; Guo et al., 2023). Following the 5-minute incubation period, the reaction was then immediately quenched by five thorough washes with PBS (phosphate buffered saline; Thermo Fisher) containing 10 mM sodium ascorbate (Spectrum Chemicals), 5 mM Trolox (Sigma-Aldrich), and 10 mM sodium azide (Sigma-Aldrich), all at room temperature. For biochemical characterization or proteomic sample collection, the quenching solution was then drained, after which brains were snap frozen in liquid nitrogen (and minimal residual quenching solution) and stored at −80°C. For immunocytochemistry, the brains were immediately fixed and stained (see below for details).

### Lysis of fly brains

Fly brains were lysed using the methods previously described by Li et al. 2020 (Cell). To avoid potential loss during transfer, brains were processed in the original sample tube. Following biotinylation, 40 μL of high-SDS RIPA buffer (50 mM Tris-HCl [pH 8.0], 150 mM NaCl, 1% sodium dodecyl sulfate (SDS), 0.5% sodium deoxycholate, 1% Triton X-100, 1x protease inhibitor cocktail (P8849), and 1 mM phenylmethylsulfonyl fluoride (PMSF); Sigma-Aldrich) was added to each sample (i.e., EP tube) of frozen brains. Disposable pestles driven by an electric motor (Thermo Fisher) were then used to extensively grind the samples on ice. Following processing, brain lysates of the same experimental group were merged into a single tube, for a final volume of 300 μL of high-SDS RIPA buffer per experimental group. Samples were then vortexed briefly, followed by two rounds of 10-second sonication (Li et al., 2020), after which they were heated to 95°C for 5 minutes to denature postsynaptic density (Li et al., 2020; Loh et al., 2016). Next, 1.2 mL of SDS-free RIPA buffer (50 mM Tris-HCl [pH 8.0], 150 mM NaCl, 0.5% sodium deoxycholate, 1% Triton X-100, 1x protease inhibitor cocktail (P8849), and 1 mM PMSF) was added to each sample (i.e., experimental group), after which they were rotated for 1 hour at 4°C. Brain lysates were then transferred to 3.5 mL ultracentrifuge tubes (Beckman Coulter), which contained 200 μL of normal RIPA buffer (50 mM Tris-HCl [pH 8.0], 150 mM NaCl, 0.2% SDS, 0.5% sodium deoxycholate, 1% Triton X-100, 1x protease inhibitor cocktail (P8849), and 1 mM PMSF). Lysates were then centrifuged at 100,000 g for 30 minutes at 4°C, after which 1.5 mL (per sample) of the supernatant was carefully collected and kept on ice.

### Enrichment of biotinylated proteins

Streptavidin magnetic beads (Pierce) were used to enrich (i.e., “capture”) biotinylated proteins from brain lysates. All chemicals were purchased through Sigma-Aldrich. For silver stains and Western blots, 20 μL of streptavidin beads were added to each sample (which each contained ∼50 dissected brains). For proteomic samples, 400 μL were added to each sample (which contained ∼1000 dissected brains). Before being added to brain lysates, streptavidin beads were washed twice with normal RIPA buffer (50 mM Tris-HCl [pH 8.0], 150 mM NaCl, 0.2% SDS, 0.5% sodium deoxycholate, and 1% Triton X-100). Washed beads were then added to the post-ultracentrifugation brain lysates and incubated on a rotator overnight at 4°C. Following incubation, beads were washed twice with 1 mL of normal RIPA buffer, then once with 1 mL of 1 M KCl, once with 1 mL of 0.1 M Na2CO3, once with 1 mL of 2 M urea in 10 mM Tris-HCl [pH 8.0], and finally twice with 1 mL of normal RIPA buffer. For silver stains and Western blots, biotinylated proteins were eluted by heating beads at 95°C for 10 minutes in 20 μL of 3x protein loading buffer (Bio-Rad), supplemented with 20 mM dithiothreitol (DTT) and 2mM biotin. For proteomic samples, on-bead trypsin digestion was performed following enrichment (see below for details).

### Streptavidin blot

Labeled proteins were separated using electrophoresis. For protein electrophoresis, 4–12% Bis-Tris PAGE gels (Thermo Fisher) were used, following the manufacturer’s protocol. Proteins were then transferred to nitrocellulose membranes (Thermo Fisher) for Streptavidin blots. All wash and incubation steps were performed on an orbital shaker at room temperature. For Streptavidin blots, samples were blocked with 3% bovine serum albumin (BSA) in TBST (Tris-buffered saline with 0.1% Tween 20; Thermo Fisher) for 1 hour. Samples were then incubated with HRP-conjugated streptavidin. Clarity Western ECL blotting substrate (Bio-Rad) and BioSpectrum imaging system (UVP) were used for detection of biotinylated proteins.

### On-bead trypsin digestion of biotinylated proteins

Proteomic samples were prepared for mass spectrometry analysis using the methods previously described by Li et al. 2020 (Cell). To remove non-biotinylated proteins while retaining the biotinylated proteins on the beads, samples were washed twice with 200 μL of 50 mM Tris-HCl buffer [pH 7.5], followed by two washes with 2 M urea/50 mM Tris [pH 7.5] buffer. Next, the final volume of 2 M urea/50 mM Tris [pH 7.5] buffer was removed, and samples were incubated with 80 μL of 2 M urea/50 mM Tris buffer containing 1 mM dithiothreitol (DTT) and 0.4 μg trypsin. Samples were then incubated in the urea/trypsin buffer for 1 hour at 25°C while shaking at 1000 revolutions per minute (rpm). After 1 hour, the supernatant was removed and transferred to a fresh tube. The streptavidin beads were washed twice with 60 μL of 2 M urea/50 mM Tris [pH 7.5] buffer and the washes were combined with the on-bead digest supernatant. The eluate was reduced with 4 mM DTT for 30 minutes at 25°C with shaking at 1000 rpm. The samples were alkylated with 10 mM iodoacetamide and incubated for 45 minutes in the dark at 25°C while shaking at 1000 rpm. An additional 0.5 μg of trypsin was added to the sample and the digestion was completed overnight at 25°C with shaking at 700 rpm. After overnight digestion, the sample was acidified (pH < 3) by adding formic acid (FA) such that the sample contained ∼1% FA. Samples were desalted on C18 StageTips (3M). Briefly, C18 StageTips were conditioned with 100 μL of 100%MeOH, 100 μL of 50%MeCN/0.1% FA, and 2x with 100 μL of 0.1% FA. Acidified peptides were loaded onto the conditioned StageTips, which were subsequently washed 2x with 100 μL of 0.1%FA. Peptides were eluted from StageTips with 50 μL of 50%MeCN/0.1% FA and dried to completion.

### TMT labeling and SCX StageTip fractionation of peptides

Desalted peptides were labeled with 8 TMT reagents from a 10-plex reagent kit (Thermo Fisher), as directed by the manufacturer. Peptides were reconstituted in 100 μL of 50 mM HEPES buffer. Each 0.8 mg vial of TMT reagent was reconstituted in 41 μL of anhydrous acetonitrile and added to the corresponding peptide sample for 1 hour at room temperature. Labeling of samples with TMT reagents was completed with the design described in Fig. 1A. TMT labeling reactions were quenched with 8 μL of 5% hydroxylamine at room temperature for 15 minutes with shaking, evaporated to dryness in a vacuum concentrator, and desalted on C18 StageTips as described above.

For the TMT 8-plex cassette, 50% of the sample was fractionated by Strong Cation Exchange (SCX) StageTips while the other 50% of each sample was reserved for LC-MS analysis by a single shot, long gradient. One SCX StageTip was prepared per sample using 3 plugs of SCX material (3M) topped with 2 plugs of C18 material. StageTips were sequentially conditioned with 100 μL of MeOH, 100 μL of 80%MeCN/0.5% acetic acid, 100 μL of 0.5% acetic acid, 100 μL of 0.5% acetic acid/500mM NH4AcO/20% MeCN, followed by another 100 μL of 0.5% acetic acid. The dried sample was re-suspended in 250 μL of 0.5% acetic acid, loaded onto the StageTips, and washed 2x with 100 μL of 0.5% acetic acid. The sample was trans-eluted from C18 material onto the SCX with 100 μL of 80%MeCN/0.5% acetic acid, and consecutively eluted using 3 buffers with increasing pH. The first elution was with pH 5.15 (50mM NH4AcO/20% MeCN), followed by pH 8.25 (50mM NH4HCO3/20% MeCN), and finally with pH 10.3 (0.1% NH4OH, 20% MeCN).

Three eluted fractions were re-suspended in 200 μL of 0.5% acetic acid, to reduce the MeCN concentration, and subsequently desalted on C18 StageTips as described above. Desalted peptides were dried to completion.

### Liquid chromatography and mass spectrometry

Desalted, TMT-labeled peptides were resuspended in 9 μL of 3% MeCN, 0.1% FA and analyzed by online nanoflow liquid chromatography tandem mass spectrometry (LC-MS/MS) using a Fusion Lumos mass spectrometer (Thermo Fisher) coupled on-line to a Proxeon Easy-nLC 1000 (Thermo Fisher). Four microliters of each sample was loaded at 500 nl/min onto a microcapillary column (360 μm outer diameter × 75 μm inner diameter) containing an integrated electrospray emitter tip (10 μm), packed to approximately 24 cm with ReproSil-Pur C18-AQ 1.9 μm beads (Dr. Maisch GmbH) and heated to 50°C. The HPLC solvent A was 3% MeCN, 0.1% FA, and the solvent B was 90% MeCN, 0.1% FA. Peptides were eluted into the mass spectrometer at a flow rate of 200 nl/min. The SCX fractions were run with 110-minute method, which used the following gradient profile: (min:%B) 0:2; 1:6; 85:30; 94:60; 95:90; 100:90; 101:50; 110:50 (the last two steps at 500 nL/min flow rate). Non-fractionated samples were analyzed using a 260 min LC-MS/MS method with the following gradient profile: (min:%B) 0:2; 1:6; 235:30; 244:60; 245:90; 250:90; 251:50; 260:50 (the last two steps at 500 nL/min flow rate). The Fusion Lumos was operated in the data-dependent mode acquiring HCD MS/MS scans (r =50,000) after each MS1 scan (r = 60,000) on the top 12 most abundant ions using an MS1 target of 3 × 106 and an MS2 target of 5 × 104. The maximum ion time utilized for MS/MS scans was 120 ms; the HCD normalized collision energy was set to 34; the dynamic exclusion time was set to 20 s, and the peptide match and isotope exclusion functions were enabled. Charge exclusion was enabled for charge states that were unassigned, 1 and >7.

### Mass spectrometry data processing

Collected data were analyzed using the Spectrum Mill software package v6.1 pre-release (Agilent Technologies). Nearby MS scans with the similar precursor m/z were merged if they were within ± 60 s retention time and ±1.4 m/z tolerance. MS/MS spectra were excluded from searching if they failed the quality filter by not having a sequence tag length 0 or did not have a precursor MH+ in the range of 750 – 4000. All extracted spectra were searched against a UniProt database containing *Drosophila melanogaster* reference proteome sequences. Search parameters included: ESIC EXACTIVE-HCD-v2 scoring parent and fragment mass tolerance of 20 ppm, 30% minimum matched peak intensity, trypsin allow P enzyme specificity with up to four missed cleavages, and calculate reversed database scores enabled. Fixed modifications were carbamidomethylation at cysteine. TMT labeling was required at lysine, but peptide N termini were allowed to be either labeled or unlabeled. Allowed variable modifications were protein N-terminal acetylation and oxidized methionine. Individual spectra were automatically assigned a confidence score using the Spectrum Mill auto-validation module. Score at the peptide mode was based on target-decoy false discovery rate (FDR) of 1%. Protein polishing auto-validation was then applied using an auto thresholding strategy. Relative abundances of proteins were determined using TMT reporter ion intensity ratios from each MS/MS spectrum and the median ratio was calculated from all MS/MS spectra contributing to a protein subgroup. Proteins identified by 2 or more distinct peptides and ratio counts were considered for the dataset.

### Proteomic data analysis

For our proteomic data analysis, we adopted the ratiometric and cutoff methodologies as previously described (Hung et al., 2014; Li et al., 2020). Detected proteins were classified according to the annotation of subcellular localization in the UniProt database. Proteins with the plasma membrane and extracellular matrix annotations were classified as true-positives (TPs). Proteins with either nucleus, mitochondrion, or cytoplasm annotation, but without the membrane annotation, were classified as false-positives (FPs). Of the total 2,007 detected proteins (those with 2 or more unique peptides), 380 were TPs and 671 were FPs. For each replicate, the proteins were first ranked in a descending order according to the TMT ratio (128C:129C, 129N:130N, 126:127C, or 127N:128N) (Fig. 1D). For each protein on the ranked list, the accumulated true-positive count and false-positive count above its TMT ratio were calculated.

To determine the optimal TMT cutoff values, we plotted the receiver operating characteristic (ROC) curve for each biological replicate (Fig. 1F; Fig. S1), which depicts the true-positive rate against the false-positive rate of detected proteins (Li et al., 2020; Nahm, 2022). To maximize the signal-to-noise ratio of the proteomes, each biological replicate was cut off at the position where the value of *true-positive rate (i.e., sensitivity) – false-positive rate (i.e., specificity)* was maximized: 128C:129C = 0.3076, 129N:130N = 0.3316, 126:127C = 0.3027, or 127N:128N = 0.3059. Post-cutoff proteomic lists of the two biological replicates for each time point were intersected to obtain the final proteome (i.e., only TP proteins that overlapped in both biological replicates at each time point were selected). We also performed cutoff analyses with a different TMT pairing regime (126:128N, 127N:127C, 128C:130N, and 129N:129C) and obtained very similar proteomes (data not shown). All ROC curves were plotted using PEELing, which is an integrated and user-centric platform for spatially-resolved proteomics data analysis (Peng et al., 2025).

To generate correlation plots (Fig. 1E), both biological replicates for each age group were compared to one control at a time, either the control lacking HRP or the control lacking H2O2. For example, for the correlation plot investigating the correlation between biological replicates for old flies using the “no HRP” control, 126:128N is plotted on the x-axis and 127N:128N is plotted on the y-axis. All correlation plots were done using the tidyverse (Wickham et al., 2019) and ggpubr (Kassambara, 2023) packages in R (R Studio Version 2024.12.0+467; R statistical software version 4.1.0; R Core Team, 2021).

### Quantitative comparison of young and old proteomes

For the cellular component Gene Ontology (GO) analysis (Fig. 1G), we uploaded the final proteomes (i.e., the 872 proteins identified following our ratiometric and cutoff analyses) into the STRING database search portal and plotted the top five retrieved terms on cellular compartment with the lowest false discovery rates (Szklarczyk et al., 2023). For the biological process GO analyses (Fig. 2B), we uploaded ∼100 of the most up-regulated and ∼100 of the most down-regulated genes into the STRING database search portal and plotted the top five retrieved terms on biological process with the lowest false discovery rates (Szklarczyk et al., 2023). For the protein network (Fig. 2C), we uploaded ∼100 of the most up-regulated and ∼100 of the most down-regulated genes into the STRING database search portal and clustered their corresponding proteins by their reported protein-protein interactions and corresponding confidence scores using a Markov clustering algorithm (inflation value set at 2.5) (Szklarczyk et al., 2023). The resulting protein-protein interactions (PPI) were then plotted in Cytoscape (v3.10.3) (Gustavsen et al., 2019). PPI plot “hub” genes were identified using the Cytoscape cytoHubba plugin (Chin et al., 2014). FlyBase and UniProt were searched to determine if each protein in the protein-protein interaction (PPI) plot had reported function in synapse organization, synapse regulation, cell adhesion, development, and localization and transport.

For the volcano plot (Fig. 2A), the TMT ratio (log2FoldChange) for each of the 872 proteins identified was plotted on the x-axis and the log10(Ratio P-value) was plotted on the y-axis using the ggplot2 (Wickham, 2016) and ggrepel (Slowikowski, 2024) packages in R (R Studio Version 2024.12.0+467; R statistical software version 4.1.0; R Core Team, 2021). To identify approximately 50 candidate genes predicted to be involved in brain aging for further screening, we generated a volcano plot, where the TMT ratio (log2FoldChange) for each of the 872 proteins identified was plotted on the x-axis and the log10(Ratio P-value) was plotted on the y-axis (Fig. 2A). Genes associated with the largest change in protein levels between time points were identified using arbitrary cutoff values: 0.4 to -0.4 log protein ratios and 0.1 p-value (Fig. 2A). This generated a list of 71 genes, which were then assessed individually to determine if they were located either on the cell-surface or in the extracellular space. Candidate genes were selected for further screening only if they were known (or predicted to be) located either on the cell-surface or extracellular space. In total, 48 candidate genes were selected for further screening, most of which were known to have at least one strong human ortholog (Supplementary Table 2) (Wang et al., 2017; Wang et al., 2019a; Wang et al., 2019b).

### Immunohistochemistry

Dissection and immunostaining of fly brains were performed according to previously described methods (Wu and Luo, 2006; Kelly et al., 2017). Briefly, the brains were dissected in 1xPBS (phosphate buffered saline; Thermo Fisher) and then fixed in 4% paraformaldehyde (Electron Microscopy Sciences) in 1xPBS with 0.015% Triton X-100 (Sigma-Aldrich) for 20 minutes on a nutator at room temperature. Fixed brains were washed with PBST (0.3% Triton X-100 in PBS) four times, each time nutating for 20-30 minutes. The brains were then blocked in 5% normal donkey serum (Jackson ImmunoResearch) or normal goat serum (BioLegend) in PBST for 1 hour at room temperature or overnight at 4°C on a nutator. Primary antibodies were diluted in the blocking solution and incubated with brains for 36–48 hours on a 4°C nutator. After washing with PBST three times, each time nutating for 20 minutes, brains were incubated with secondary antibodies diluted in the blocking solution and nutated in the dark for 24–36 hours at 4°C. Brains were then washed again with PBST three times, each time nutating for 20 minutes. Immunostained brains were mounted with SlowFade antifade reagent (Thermo Fisher) and stored at 4°C before imaging.

Primary antibodies used in immunostaining include: rat anti-NCad (1:40; DN-Ex#8, Developmental Studies Hybridoma Bank), chicken anti-GFP (1:1000; GFP-1020, Aves Labs), rabbit anti-DsRed (1:200; 632496, Takara Bio), and mouse anti-Repo (1:100; 8D12 anti-Repo, Developmental Studies Hybridoma Bank). Donkey or goat secondary antibodies conjugated to Alexa Fluor 488/568/647 (Jackson ImmunoResearch or Thermo Fisher) were used at 1:250. Neutravidin (Thermo Fisher) pre-conjugated with Alexa Fluor 647 was used to detect biotin.

### Image acquisition and processing

Images were acquired by either a Zeiss LSM 780 laser-scanning confocal microscope (with a 40x/1.4 Plan-Apochromat oil objective, Carl Zeiss) or a Leica STELLARIS 5 confocal microscope (with a 20x objective). Confocal z-stacks were obtained by 1-μm intervals at the resolution of 512×512. Images were exported as maximum projections or single confocal sections by ZEN (Carl Zeiss) in the format of TIFF. Z-stack images were processed using Image J (Schneider et al., 2012) and Fiji (Schindelin et al., 2012) (Image-Stacks-Z projection-Max Intensity). BioRender was used to make diagrams. Adobe Illustrator was used to assemble all figures, as well as rotate and crop images.

### Fly rearing (for lifespan, climbing assays, and single-nucleus RNA-sequencing)

To rear flies for lifespan assays, climbing assays, and single-cell RNA sequencing, we placed 20-30 unmated females with 5-10 males of each genotype in bottles containing standard cornmeal medium. Flies were allowed to mate and lay eggs for 3 days, after which they were flipped into new food bottles and allowed to mate and lay eggs for an additional 4 days. On day 4, all F0 parent flies were cleared from the bottles. In general, the glial-GS was used as the maternal line, whereas the RNAi/UAS/flySAM/etc. lines were used as the paternal F0 lines. Following eclosion, F1 flies were allowed to mate for 1-3 days, after which they collected under light CO2 anesthesia and transferred into “cages” containing vials with standard cornmeal medium. Cages consisted of clear polystyrene containers (53 mm diameter, 100 mm height, 175 mL) which were plugged with ceaprene foam stoppers (for 175 mL containers) (Greiner Bio-One) that had been modified to accommodate a standard fly food vial (VWR) (i.e., a razor was used to create a hole in the middle of the stopper, which securely held a vial containing standard fly medium). Fly food vials were replaced with new food vials every 2-3 days.

In most cases, we were able to collect enough F1 flies 1-3 days post-eclosion (i.e., flies ranged from 1-3 days old when collected and transferred into the lifespan cages) from two crosses per genotype. However, in some cases certain crosses produced fewer F1 progeny, which eclosed slowly over a broader range of time. In these cases, flies could range from 1-5 days old when collected and transferred into the lifespan cages. To best control for this variability in eclosion time, equal numbers of flies were collected for each condition (Ru846 treatment vs. ethanol control) at each collection time point. For example, if 50 F1 flies were collected from a cross on Day 1 following eclosion, 25 flies would be assigned the Ru486 treatment and 25 flies would be assigned to the ethanol control. In cases where we were unable to collect sufficient progeny after 5 days, or wanted to further investigate a particular candidate (like DIP-beta), crosses were remade (with 3-5 crosses per genotype) to reduce the amount of time for sufficient numbers of F1 progeny to eclose. This approach allowed us to maintain a reasonable amount of efficiency screening a large number of candidate genes.

### Lifespan assays

For the broadscale lifespan screen of our candidate genes, we collected 80-100 F1 males and 80-100 F1 females per genotype/treatment (i.e., for F1 males, 80-100 flies were assigned to the Ru486 treatment and 80-100 flies were assigned to the ethanol control) and placed them into lifespan cages. Males and females were housed separately for the full duration of all lifespan experiments. Depending on genotype fecundity and eclosion time, flies ranged from 1-5 days old at the start of the experiment (i.e., when they were collected and placed into the lifespan cages following eclosion). All lifespan cages were kept in a humidified, temperature controlled incubator at 25℃ with a 12:12 hour light/dark cycle for the full duration of the experiment (Bauer et al., 2004). Flies were allowed to mature for 3-6 days on plain standard cornmeal medium (to ensure that the adult brains were fully developed prior to gene expression manipulation), after which flies were switched to fly food vials containing either the Ru486 (mifepristone) drug treatment or the ethanol control. To prepare the Ru486 food vials, 100 μL of Ru486 solution (4mM Ru486 powder dissolved by vortex into 200-proof ethanol) was added to each vial, which contained standard fly medium (Goodman et al., 2021). To prepare the ethanol control vials, 100 μL of 200-proof ethanol was added to each vial. The Ru486 was purchased from Cayman Chemical and the 200-proof ethanol was purchased from Thermo Scientific. All Ru486 and ethanol control food vials were allowed to dry completely before being introduced to the flies to prevent flies from sticking to the wet medium. Ru486 or ethanol control food vials were exchanged with new food vials every 2-3 days for the full duration of the experiment, at which time the number of flies that were alive, dead, or had escaped during the food exchange were noted. Lifespan experiments continued until all flies were dead.

### Climbing assays

To age flies for climbing assays (and single-cell RNA sequencing), we collected 56 F1 males and 66-68 F1 females (collected from several glial-GeneSwitch x DIP-β flySAM2.0 crosses) per treatment (Ru486 treatment vs. ethanol control) and transferred them into lifespan cages.

Flies were allowed to mature for 3-6 days on plain standard cornmeal medium, after which flies were switched to fly food vials containing either the Ru486 drug treatment or the ethanol control. Males and females were aged separately in their respective cages until they reached the target age of 50-55 days old. To prepare for climbing assays, flies were collected under light CO2 anesthesia and transferred from their “cages” into either Ru486 drug supplemented or ethanol control food vials (8-10 males and 8-10 females per vial) (Ali et al., 2011). Flies were allowed to recover from the CO2 anesthesia for 3 days (Ali et al., 2011; Burns et al., 2020; Burns and Saltz, 2024). During this time, flies were kept in a humidified, temperature controlled incubator at 25℃ with a 12:12 hour light/dark cycle (Ali et al., 2011). On the third day, flies were transferred to a humidified, temperature controlled room (25℃) (Goodman et al., 2019) and allowed to acclimate for 1 hour prior. Following this acclimation period, unanesthetized flies were gently transferred from food vials into the lanes of the climbing apparatus, which were plugged with cellulose acetate plugs (VWR) and additionally secured with general purpose laboratory tape (VWR), after which they were allowed to acclimate in their lanes for 5 minutes. For the climbing assay, flies were tapped to the bottom of an empty vial and immediately video recorded for 10 seconds (Ali et al., 2011; Goodman et al., 2019). Each test group was subjected to 3 consecutive trials with a 1 minute recovery between trials (Ali et al., 2011; Goodman et al., 2019). For analysis, the climbing height (in centimeters) for each fly at the 10-second time point was recorded. Mean climbing height values from the 3 trials per test group were used to account for technical variability (Ali et al., 2011; Goodman et al., 2019). Upon completion of the climbing assays, flies were humanely sacrificed by being placed in a freezer for a minimum of 24 hours (Burns and Saltz, 2024).

### Single-nucleus RNA-sequencing

The heads of female flies were collected at day 50 (i.e., when the flies were 50 days old), placed into 1.5 mL Eppendorf tubes, flash-frozen in liquid nitrogen, and stored at -80C. In total, two samples (each containing 20 heads) were prepared for single-nucleus RNA sequencing: one where DIP-beta had been overexpressed in glial cells from day 5-50 and one where DIP-beta expression had not been manipulated (i.e., the flies received an ethanol control instead of the Ru486 drug used to induce overexpression of DIP-beta). Single-nucleus suspensions were prepared as described in the Fly Cell Atlas (Li et al., 2022). After nuclei dissociation, the nuclei were purified from debris using the SONY SH800 FACS sorter. Nuclei were stained with Hoechst-33342 (1:1000, >5 minutes). It is typical to observe separate cell populations during FACS since some cells have variable sizes of nuclei and ploidy numbers. All populations were included to ensure every cell would be captured from the samples. Nuclei were collected in a 1.5 mL Eppendorf tube containing 200uL 1X PBS with 0.5% BSA (RNase inhibitor added). For each of the 10X Genomics runs, 250-300k nuclei were collected. The nuclei were centrifuged for 10 minutes at 1000g in 4C. conditions and then resuspended using 40uL of 1X PBS with 0.5% BSA (RNase inhibitor added). 2uL of the nuclei suspension were allocated for counting under a hemocytometer to calculate the nuclei concentration. For the high-throughput single-cell sequencing kit from 10X Genomics, we loaded 50k cells into the Chromium Controller to target a final number of 20-30k nuclei.

Single-nucleus RNA-seq libraries were prepared using the 10X Genomics Chromium Single Cell 3’ HT v3.1 protocol. All PCR reactions were performed using the BioRad C1000 Touch Thermal Cycler with the 96-Deep Well Reaction Module. We used 11 cycles for the cDNA amplification and then we used 13 cycles for the sample index PCR. As per the 10X Genomics protocol, 1:10 dilutions of amplified cDNA and the final gene expression libraries were evaluated on the Agilent Tapestation Bioanalyzer. Each library was diluted to approximately 4nM and equal volumes of each library were pooled in a NovaSeq PE150 S4 sequencing run. Pools were sequenced with the dual index configuration Read 1 28 cycles, Index 1 (i7) 10 cycles, Index 2 (i5) 10 cycles, and Read 2 90 cycles. The sequencing depth was about 30-40k reads per nucleus.

### Preprocessing and quality control of single-nucleus RNA-sequencing data

The Drosophila melanogaster genome (FlyBase Release 6.31) was downloaded along with a pre-mRNA GTF containing both intronic and exonic annotations, then indexed using Cell Ranger mkref v7.0.1 (10x Genomics) to enable accurate mapping of nucleus-derived transcripts (Zheng et al., 2017; Thurmond et al., 2019). Paired-end FASTQ reads were aligned to this custom reference using Cell Ranger count v7.0.1, producing raw feature-barcode matrices and QC metrics (Zheng et al., 2017).

To remove background (“ambient”) RNA contamination inherent to droplet-based assays and technical artifacts, we applied CellBender remove-background v0.3.2, which employs a deep generative model to distinguish true cell-derived signal from ambient noise (Fleming et al., 2023). Barcodes flagged as potential doublets were then identified and removed with scDblFinder v1.22.0, a Bioconductor package that simulates artificial doublets and uses neighborhood classification to assign doublet probabilities (Germain et al., 2021).

We next filtered nuclei based on gene and UMI counts: cells with fewer than 200 detected genes or fewer than 300 UMIs were discarded to exclude low-quality captures, while those exceeding five median absolute deviations above the median for either metric were deemed outliers and removed (Luecken and Theis, 2019). Finally, nuclei with >10 % mitochondrial UMIs—indicative of damaged or apoptotic cells—were excluded from further analysis (Ilicic et al., 2016). After all the above quality control processing, 26,261 nuclei in the –RU condition and 22,698 nuclei in the +RU condition were retained.

### Single-nucleus RNA clustering and annotation

The Ru486+ (+RU) and Ru486- (-RU, i.e., the ethanol control) matrices were merged into a single AnnData object and log-normalized. We then computed 150 principal components (PCs) to capture major sources of biological variation. To correct batch effects between conditions while preserving true transcriptomic differences, Harmony v1.0 was applied to the PC embeddings (Korsunsky et al., 2019).

Cell-type labels were transferred from the AD-FCA (specifically the whole-head data, comprising 108 fine-grained cell types) using machine learning basedlabel-transfer workflow (Lu et al., 2023; Stuart et al., 2019). A k-nearest-neighbor graph (k = 20) was built on the Harmony-corrected PCs, and the Leiden algorithm (Traag and Waltman, 2019) at resolution 6 was used to detect communities, yielding 130 clusters with high modularity and guaranteed connectivity (Traag et al. 2019). Clusters were then inspected and, where necessary, manually corrected based on marker expression and neighborhood structure, and subsequently consolidated into nine broad cell-type categories for downstream analyses.

### snRNA-seq: cell composition

To quantify how DIP-β overexpression alters cellular makeup, we counted the nuclei assigned to each broad cell type in DIP-β overexpression versus control samples. Cell-type proportions were normalized by the total number of nuclei per genotype, allowing direct comparison of enrichment or depletion across conditions.

### snRNA-seq: DEG analysis

Within each broad cell type, we identified differentially expressed genes (DEGs) between DIP-β overexpression and control samples using the Wilcoxon rank-sum test implemented in Scanpy’s rank_genes_groups function (Wolf et al., 2018), applying Benjamini–Hochberg correction for multiple testing. Genes with a false discovery rate (FDR) < 0.05 were considered significant.

### snRNA-seq: KEGG pathway analysis

Significant DEGs (|log₂ fold change| > 0.5, FDR < 0.05) were subjected to pathway enrichment analysis using the clusterProfiler R package (Yu et al., 2012), mapping to the Kyoto Encyclopedia of Genes and Genomes (KEGG) database (Kanehisa and Goto, 2000). Enriched pathways with adjusted p-values (FDR) < 0.05 were reported.

### snRNA-seq: cell-cell communication analysis

Cell-cell communication analysis was performed using our recently developed FlyPhoneDB2 (Qadiri et al., 2025). The core algorithm infers tissue-specific signaling events between cell types by calculating cell-cell interaction scores based on curated ligand-receptor pairs.

## Quantification and Statistical Analyses

No statistical methods were used to determine sample sizes, but our sample sizes were similar to those generally employed in the field (Li et al., 2020). Any brains damaged in dissection were excluded from analysis. Excel (Microsoft), Prism (GraphPad), and R (R Studio Version 2024.12.0+467; R statistical software version 4.1.0; R Core Team, 2021) were used for data analysis and plotting. Statistical methods used are described in the figure legend of each relevant panel.

## Data and Code Availability

The original mass spectra and the protein sequence databases used for searches have been deposited in the public proteomics repository MassIVE (http://massive.ucsd.edu) and are accessible at ftp://MSV000099083@massive-ftp.ucsd.edu. These datasets will be made public upon acceptance of the manuscript. Original proteomic data prior to analyses is provided in the Supplementary Table 1. snRNA-seq data has been deposited to NCBI Gene Expression Omnibus (GSE308135).

---

**Figure S1: Glial cell-surface proteomics**

**A:** We searched the 872 proteins included in our proteomic profiles against three databases: FlyBase, UniProt, and the Gene Ontology Data Archive (DOI 10.5281/zenodo.1205166; Version 2023-05-10; Ashburner et al., 2000) to determine which proteins had previous plasma membrane or extracellular matrix annotations. We found the 682 out of the 872 proteins could be validated by at least one database, further validating the quality of our cell-surface proteomic labeling.

**B:** We plotted the receiver operating characteristic (ROC) curve for each biological replicate answer observed high specificity.

**C:** We identified the top 10 “hub” genes in our PPI network and found that of the top 10 hub genes identified, 7 had reported function in cell adhesion (klg, kst) and synaptic wiring (DIP-β, DIP-δ, DIP-$, DIP-κ, DIP-ɑ), while the remaining three had reported function in neuron projection (CG33543), axon guidance (CG34353), and dendrite guidance (Toll-6).

**Figure S2: Functional screening of candidate genes.**

**A-C:** In our initial functional lifespan screen, glial overexpression of DIP-β was found to significantly extend lifespan for both male and female flies, compared to the negative controls (only the luciferase RNAi and flySAM controls are pictured here).

**Figure S3-S6: Functional screening of up-regulated candidate genes.**

**A-E:** We selected 19 up-regulated candidate genes for functional screening, which we selectively knocked down in glia to examine their influence on lifespan. For each up-regulated gene, we screened two RNAi lines (to control for possible off target effects). Of the 19 up-regulated candidates screened, we found that RNAi knockdown of 4 genes led to consistent lifespan decreases for males. Alternatively, for female flies, RNAi knockdown of 3 genes led to consistent lifespan increases.

**Figure S7-S8: Functional screening of down-regulated candidate genes.**

**A-J:** We selected 29 down-regulated candidate genes for functional screening, which we selectively overexpressed in glia to examine their influence on lifespan. For each of the 29 down-regulated genes, we screened one overexpression line (using either UAS-cDNA lines or flySAM2.0 lines), when available. In total, we were able to acquire and screen lines for 19 genes. Of these candidates, overexpression of 2 genes led to lifespan increases for males, whereas overexpression of 3 genes led to lifespan decreases. Alternatively, overexpression of 10 genes led to lifespan increases for females, whereas overexpression of 1 gene led to decreased lifespan. We found that constitutive overexpression of DIP-β in glia significantly increased lifespan in both males and females compared to controls.

**Figure S9: Single-nucleus RNA-sequencing - investigating the downstream transcriptomic effects of pan-glial DIP-β overexpression.A:** In total, we sequenced approximately 49,000 cells from our fly head samples, with approximately 23,000 from the glial DIP-β overexpression, (i.e., Ru486+) sample and approximately 26,000 from the negative control (i.e., Ru486-) sample.

**B:** Detailed cell subtype cluster composition annotations.

**C:** A detailed analysis investigating differentially expressed genes (DEGs) across cell subtypes.

**Supplementary Table 1: Original proteomic data prior to analyses.**

**Supplementary Table 2: Candidate genes selected for further investigation.**

**a:** In total, we selected 29 down-regulated candidate genes for functional screening. Candidate genes were selected for further screening only if they were known (or predicted) to be located either on the cell-surface or in the extracellular space.

**b:** In total, we selected 19 up-regulated candidate genes for functional screening. Candidate genes were selected for further screening only if they were known (or predicted) to be located either on the cell-surface or extracellular space. Most candidate genes selected for further screening were found to have at least one strong human ortholog (Wang et al., 2017; Wang et al., 2019a; Wang et al., 2019b).

**Supplementary Table 3: RNAi lines used to screen up-regulated candidate genes and OE lines used to screen down-regulated candidate genes.**

For each of the 19 up-regulated candidate genes, we screened 2 RNAi transgenic lines to control for potential off-target effects. For each of the 29 down-regulated genes, we screened one overexpression line (using either UAS-cDNA lines or flySAM2.0 lines), when available. Of the 29 down-regulated genes selected, we were able to acquire and screen lines for 19 genes.

