## Supplementary figures and images for "*In-situ* glial cell-surface proteomics identifies pro-longevity factors in *Drosophila*"

### supplemental figures

#### Fig. S1

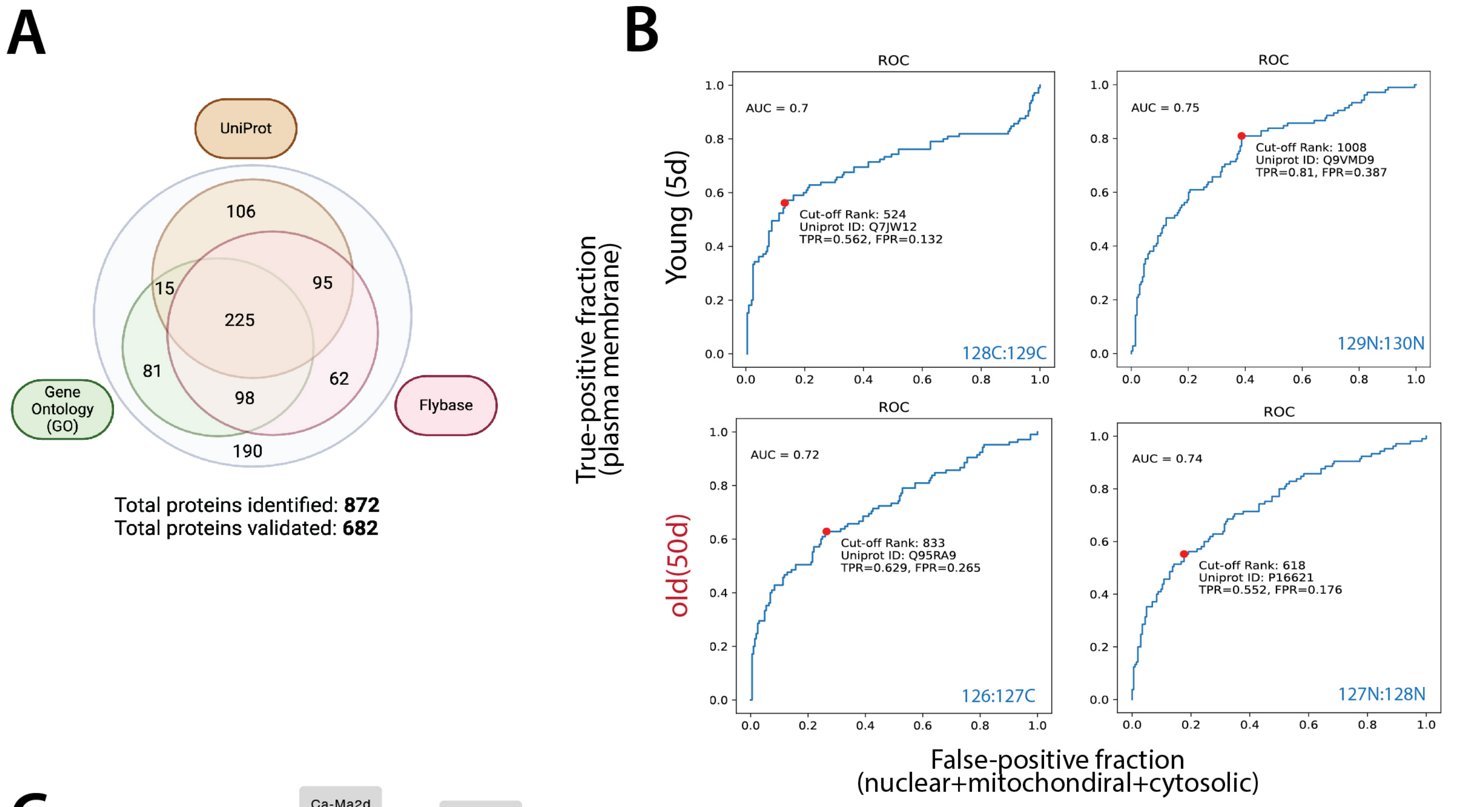


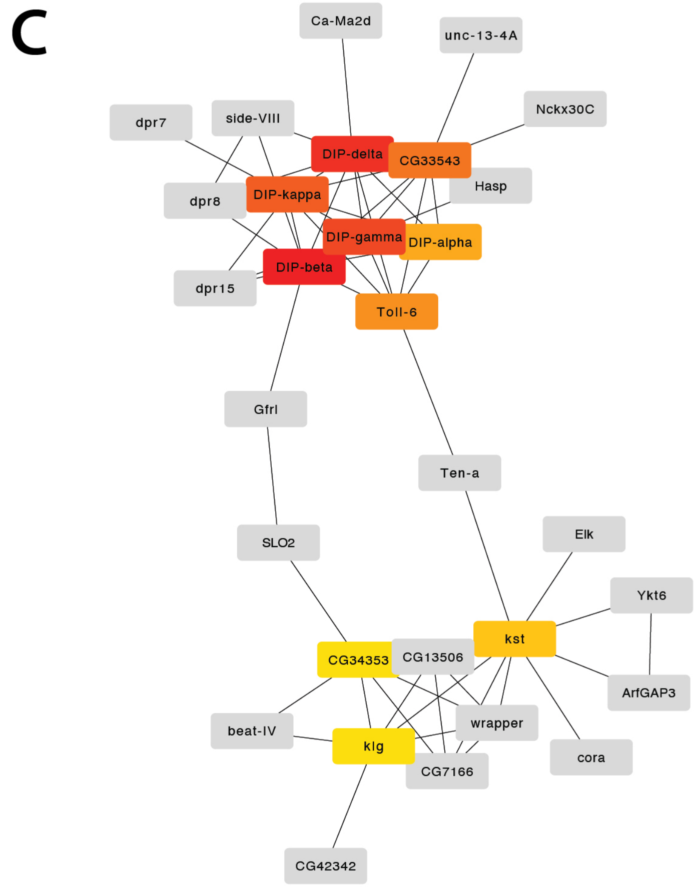


| Hub Gene | Rank |
|---|---|
| DIP-beta | 1 |
| DIP-delta | 2 |
| DIP-gamma | 3 |
| DIP-kappa | 4 |
| CG33543 | 5 |
| Toll-6 | 6 |
| DIP-alpha | 7 |
| kst | 8 |
| klg | 9 |
| CG34353 | 9 |

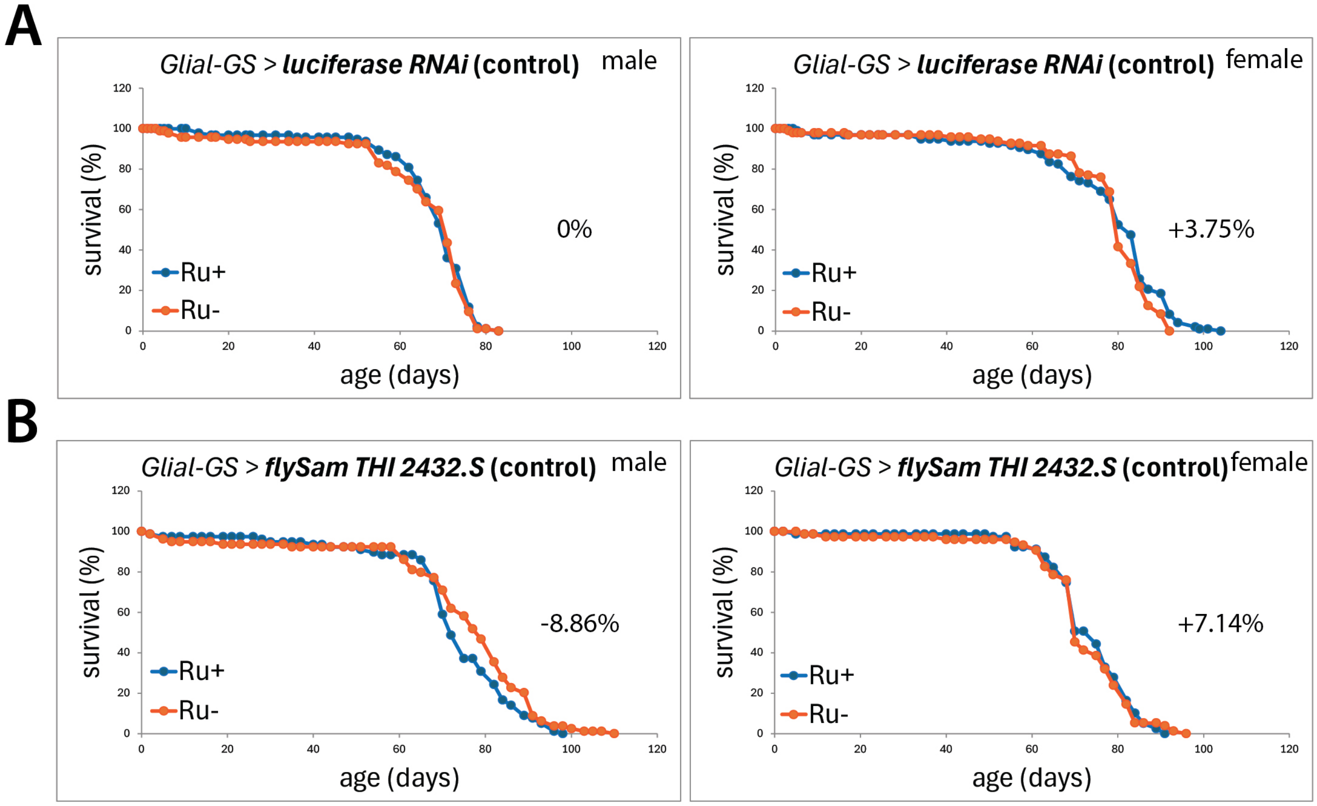

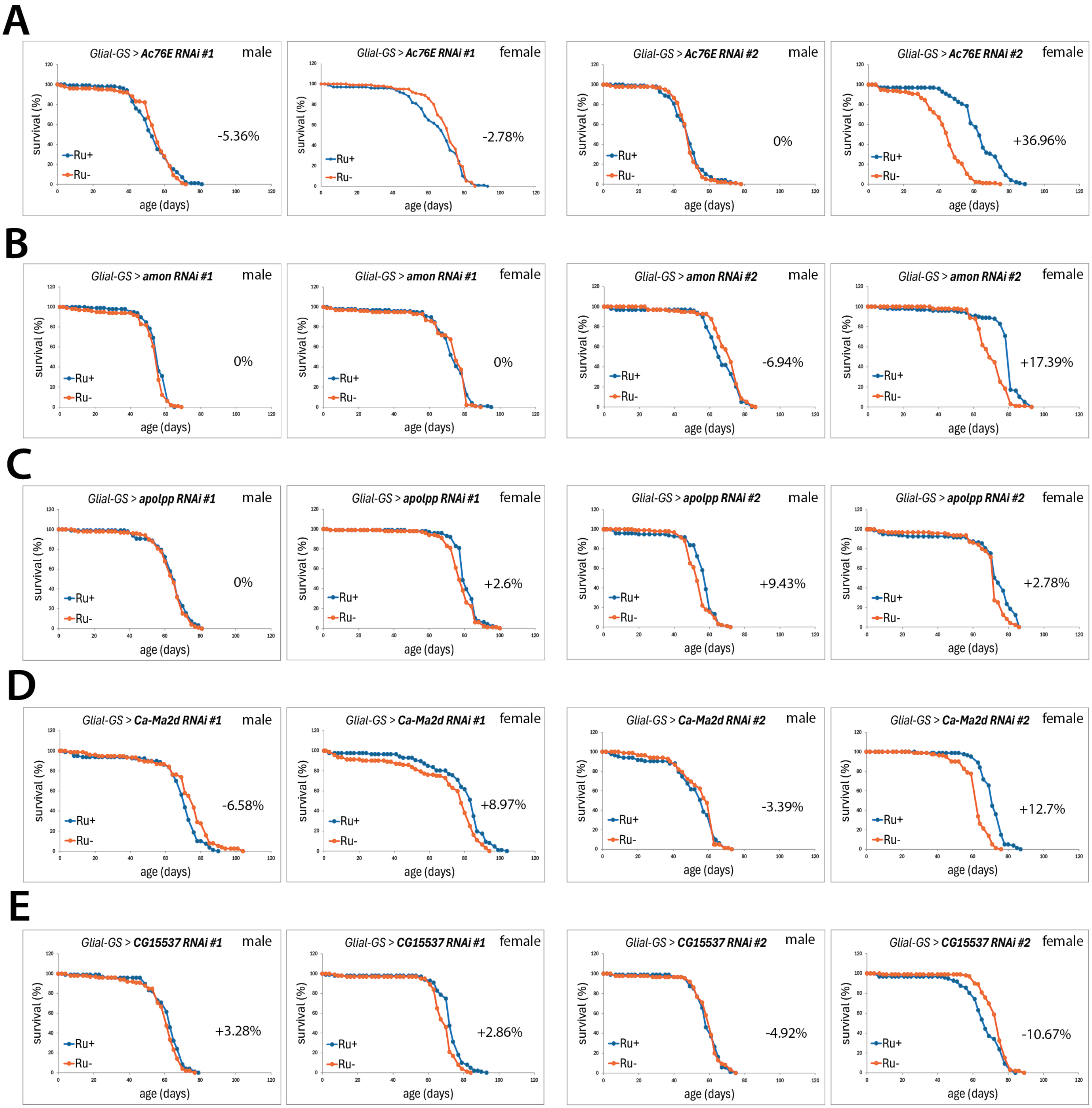

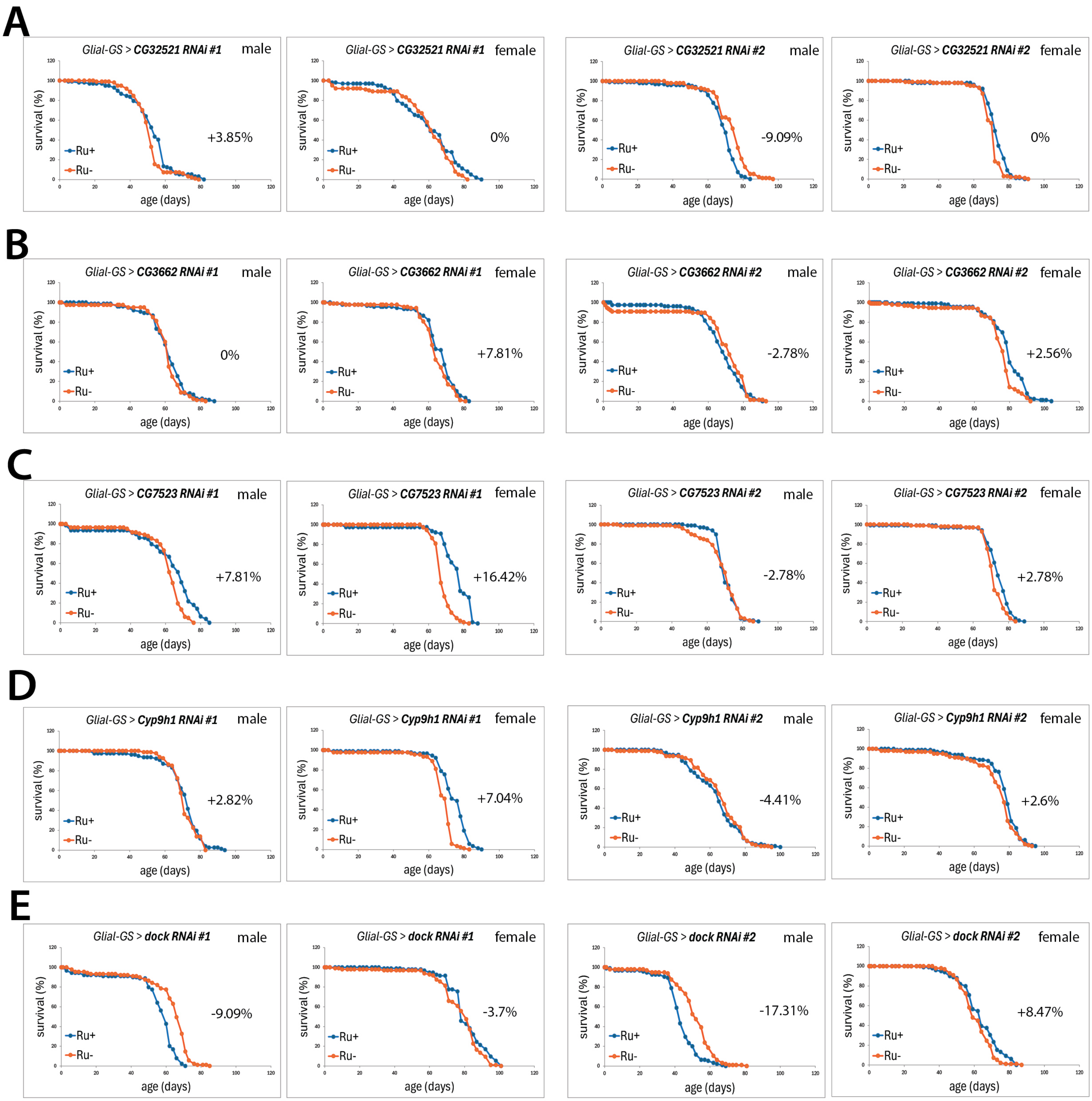

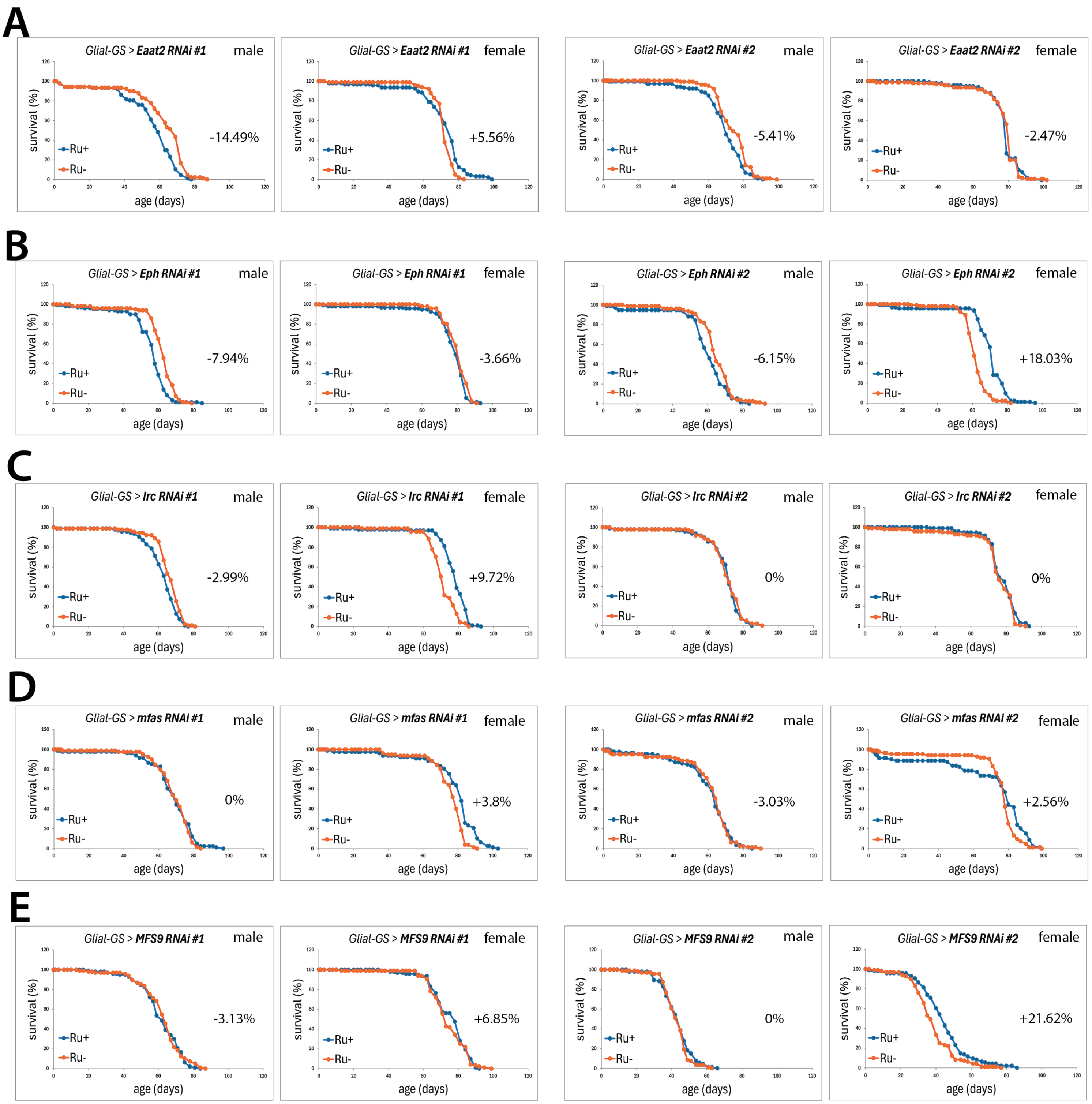

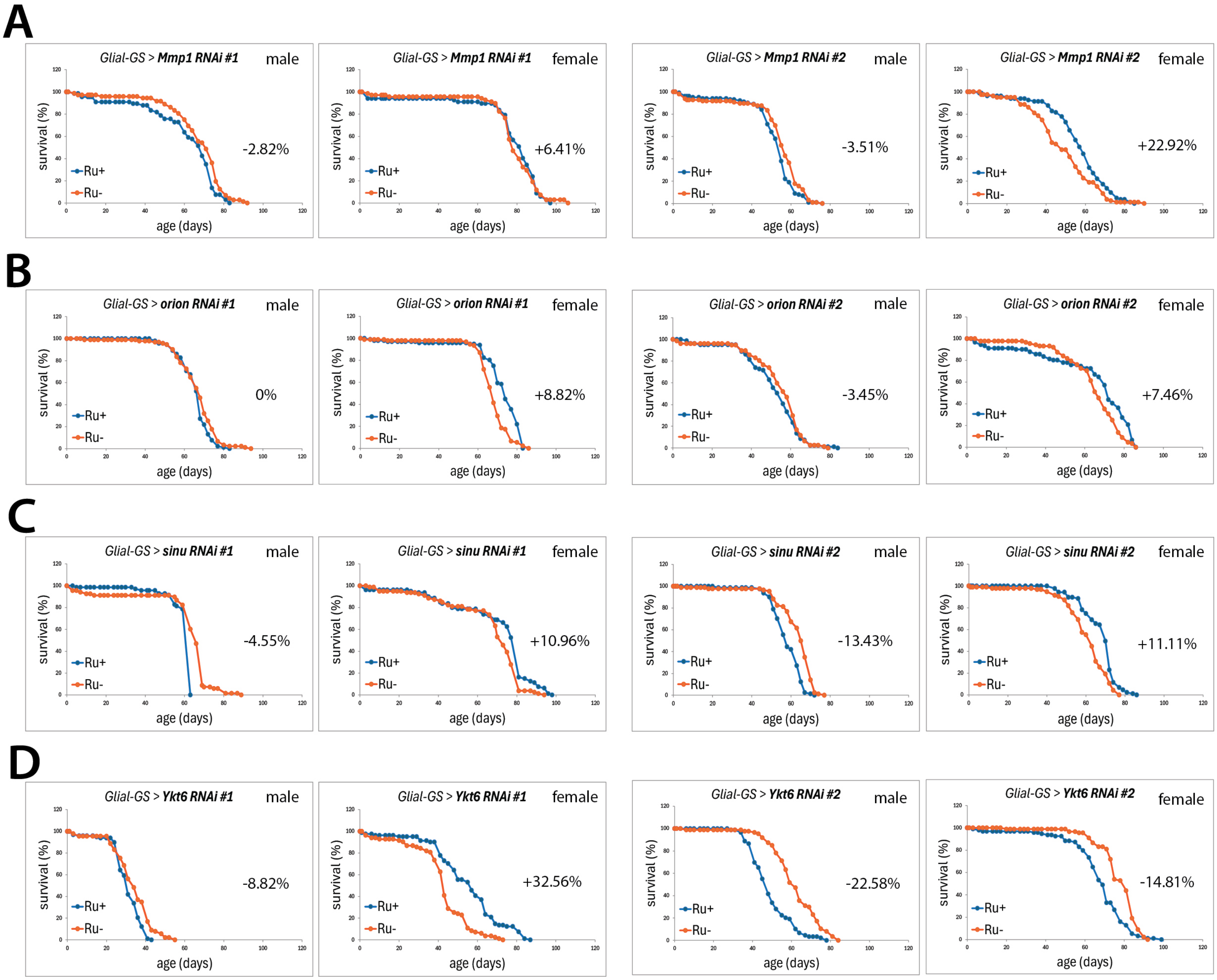

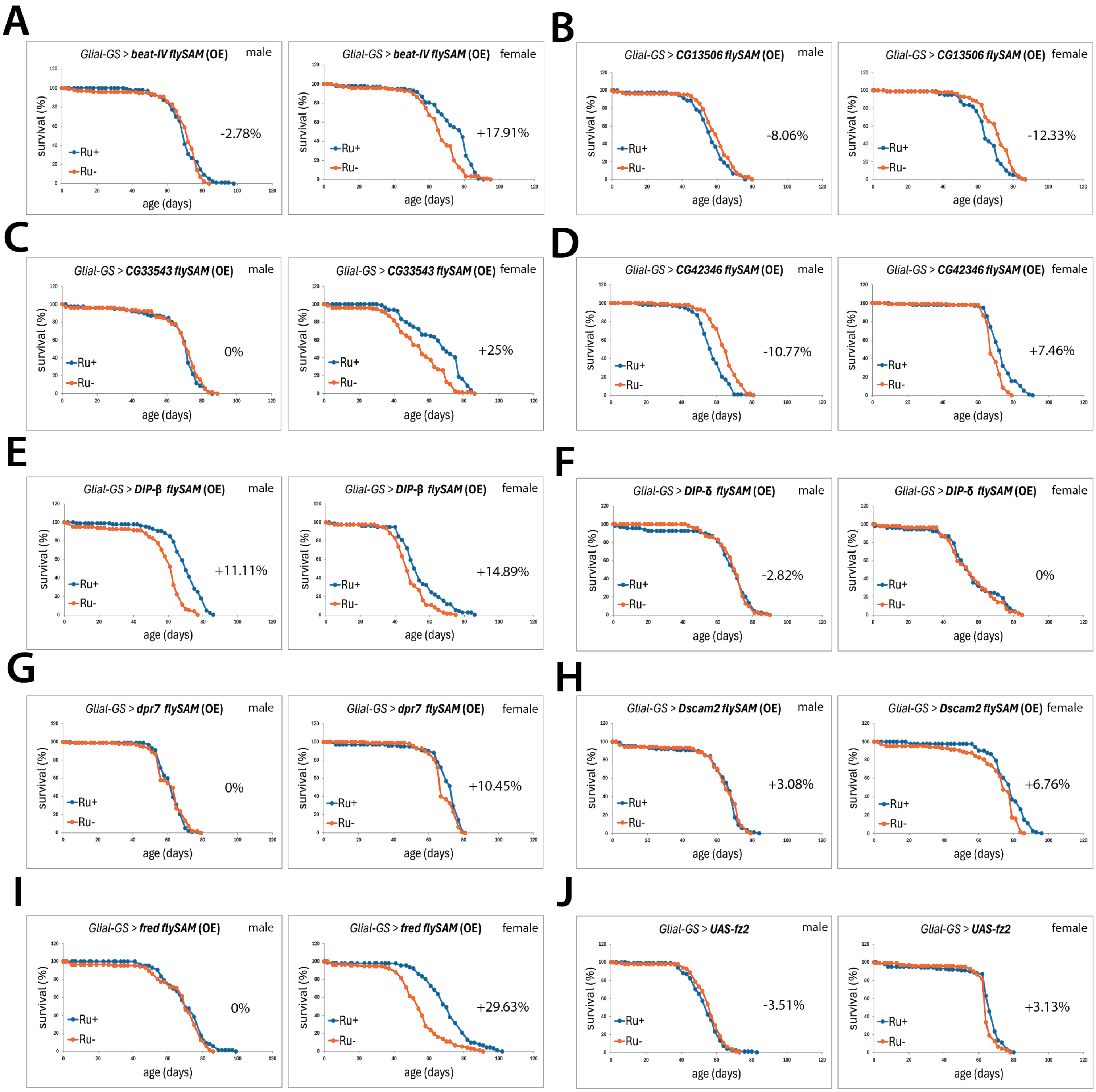

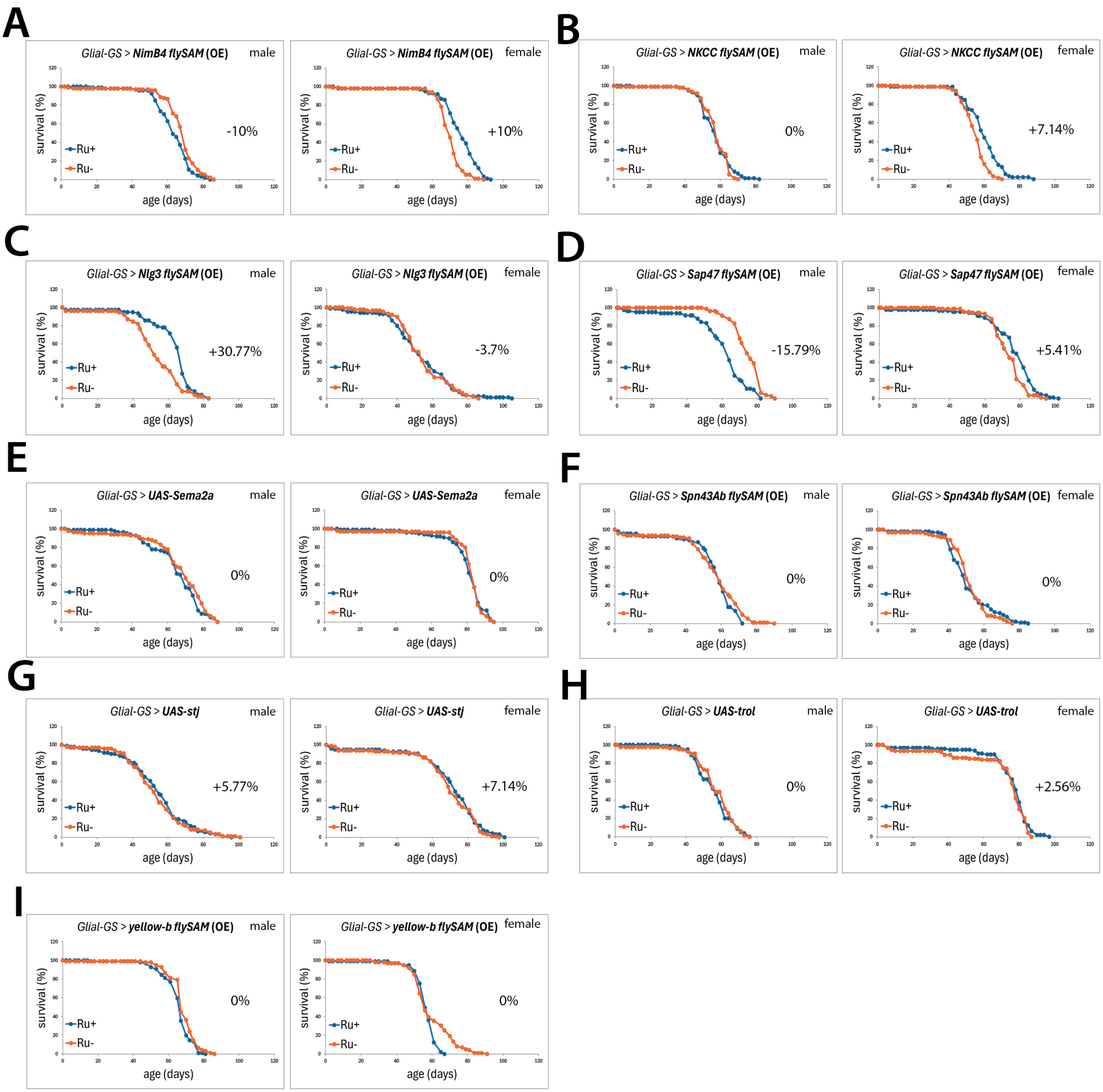

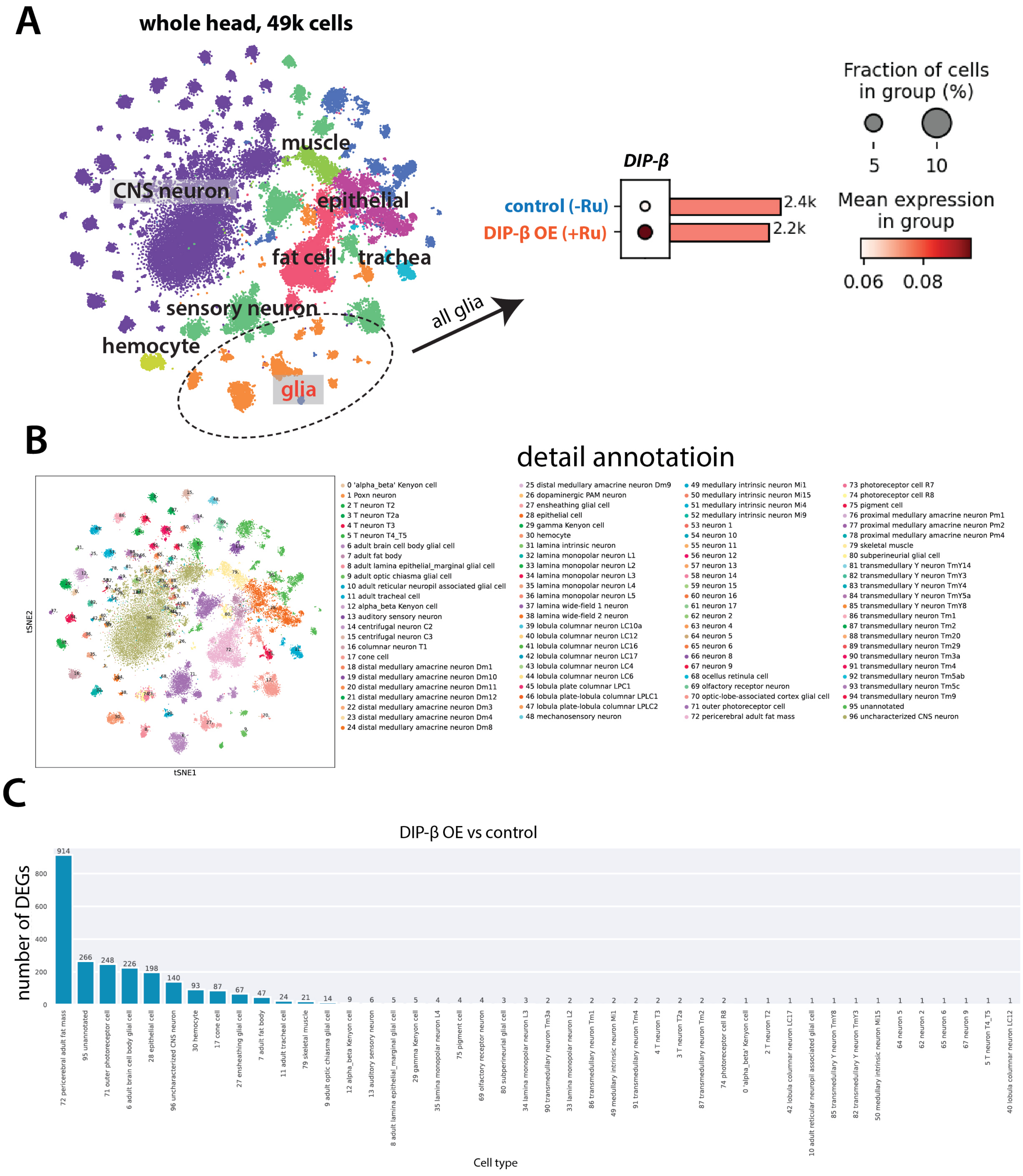
